# Transglutaminase 2 predicts parasitic worm-mediated protection against hepatic steatosis in obese mice

**DOI:** 10.1101/2025.11.21.689654

**Authors:** Samson Folami, Mona Sharma, Rugivan Sabaratnam, Guillaume Bidault, Marta Diaz-del Castillo, Martin Dale, Laura Jessica Myhill, Kristoffer Jensen Kolnes, Vaishnavi Misra, Claus Bogh Juhl, Antonio Vidal-Puig, Kurt Højlund, Johan Palmfeldt, Andrew Richard Williams, Peter Nejsum, Vineesh Indira Chandran

## Abstract

Parasitic worm infection can mitigate high fat diet-induced chronic inflammation in obese mice by reducing hepatic fat accumulation and improving insulin sensitivity. However, the molecular alterations during infection-mediated regulation of hepatic steatotic events in obesity remain poorly understood. Here, we integrated proteomic and metabolomic analyses of infected obese mice, a functional deworming and interleukin-4c (IL-4c) treated animal model, and an obese patient cohort before and after bariatric surgery to uncover molecular targets/pathways indicative of the regulation of hepatic steatosis by gut *Heligmosomoides polygyrus bakeri* (*H. p. bakeri*) helminth infection. Proteomic analysis identified alterations in several molecules related to metabolism and showed elevated levels of transglutaminase 2 (TGM2) proportionate with an infection-regulated lipid load in diet-induced obese (DIO) mice. The role of TGM2 as a reliable predictor of hepatic steatosis was validated using deworming experiments which revealed a decrease in the liver tissue levels of *Tgm2* following removal of *H. p. bakeri*. Further validation showing increase in *Tgm2* levels in IL-4c treated mice confirmed its relevance as a general response indicator of type 2 immune response induced by helminth infection. Assessment of circulating TGM2 levels in matched obese patients before and after bariatric surgery revealed no change, indicating that differential expression of TGM2 is a unique functional response to regulation of hepatic steatosis by *H. p. bakeri* infection. Together, our data show that TGM2 is a unique predictive marker of improvement in hepatic lipid profile and provides additional evidence that parasitic worm infection protects against hepatic steatosis in DIO mice. These findings pave the way for novel therapeutic opportunities to resolve hepatic steatosis and prevent its progression to severe forms of metabolic dysfunction-associated steatotic liver disease, based on TGM2 coupled with mimetics of *H. p. bakeri*-derived products.

**Graphical abstract:** 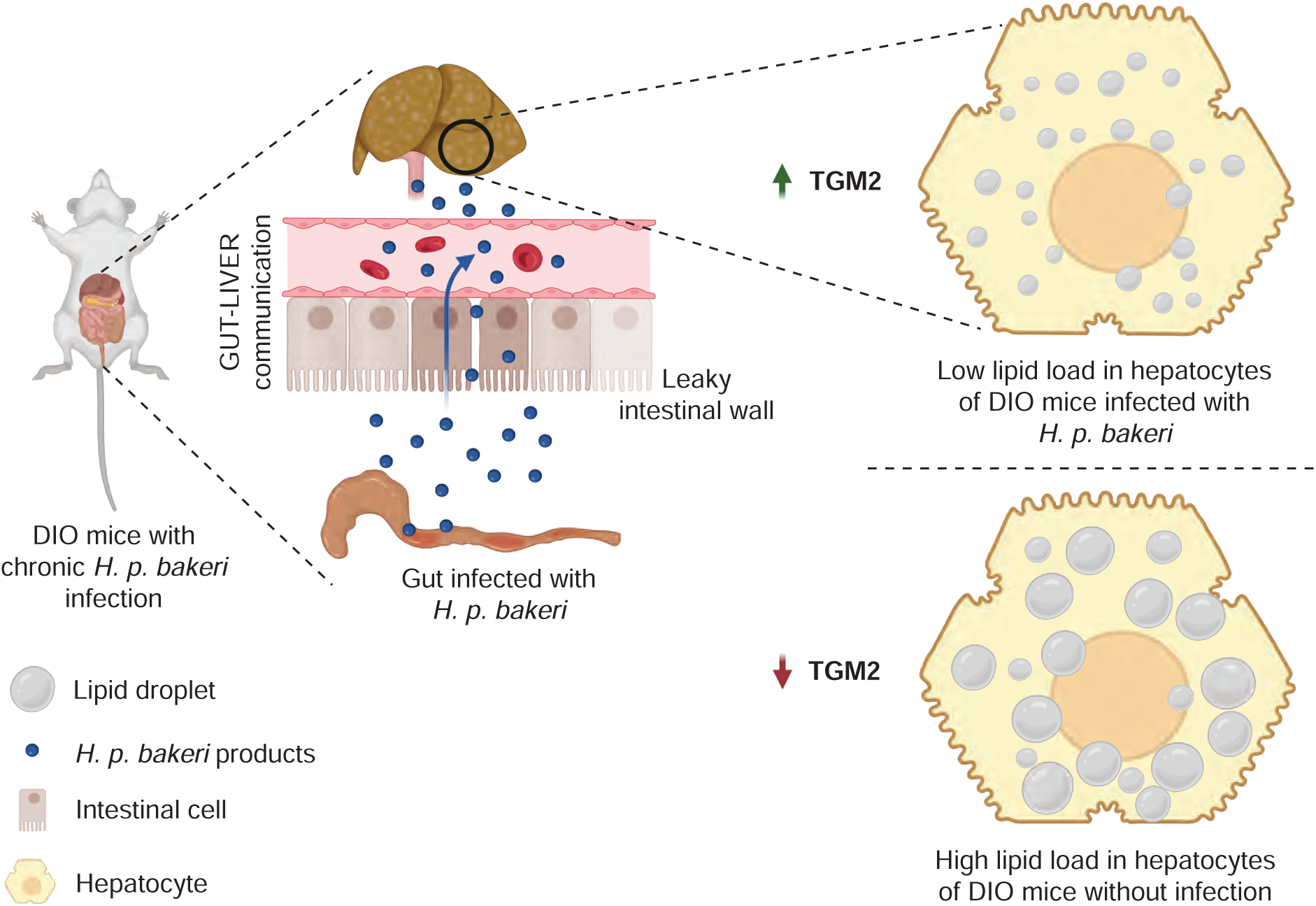

## Introduction

Parasitic worm (helminth) infections affect approximately 24% of the world’s population, especially in countries of the global south [1]. Epidemiological studies have revealed an inverse correlation between the prevalence of obesity-associated metabolic diseases and exposure to parasitic helminths, a phenomenon popularly referred to as the “hygiene hypothesis” [2, 3]. This hypothesis indicated the beneficial effects of helminth infections and their potential therapeutic properties on the host [4, 5]. In fact, studies in diet-induced obese (DIO) mice found that *Heligmosomoides polygyrus bakeri* (*H. p. bakeri)* helminth infection protects against obesity-associated metabolic complications [6]. *H. p. bakeri* is a natural parasite of mice that causes chronic infection in the gut with potent immunomodulatory effects [7]. Further investigations revealed that *H. p. bakeri* infection induces a strong modified type 2 (Th2)/Treg immune response in the gut, which is mainly mediated by interleukin-4 (IL-4) cytokine and is critically important for controlling obesity and infection by expelling the worm [8, 9]. More specifically, *H. p. bakeri* infection improved insulin sensitivity, reduced liver fat accumulation, and mitigated obesity-related chronic inflammation in a high fat DIO mouse model [10]. Intriguingly, parasitic worm infection has been shown to regulate metabolic pathologies through excretory/secretory products (ESPs), such as extracellular vesicles, glycans, microRNAs, and proteins [4, 11–15]. However, there is currently limited understanding of the molecular intermediates in the liver of DIO mice that are perturbed during *H. p. bakeri* infection-mediated regulation of steatosis/inflammatory events. A greater understanding could enable the development of mimetics of *H. p. bakeri* products or other therapeutic modalities for early intervention in fatty liver disease that can arrest its progression to more severe forms such as metabolic dysfunction-associated steatotic liver disease (MASLD)/Metabolic dysfunction-associated steatohepatitis (MASH).

In this study, we performed untargeted proteomics and metabolomics in infected DIO mice, combined them with deworming and IL-4c-treated animal models, and evaluated matching obese patients before and after bariatric surgery to uncover molecular targets/pathways involved in the regulation of hepatic steatosis by intestinal *H. p. bakeri* infection. We identified transglutaminase 2 (TGM2) as a novel indicator of infection-induced changes in lipid load in hepatocytes. Experiments in the deworming and IL-4c-treated animal models validated the changes in TGM2 levels as a unique functional response to parasitic worm infection, which were further confirmed in a matched obese patient cohort before and after bariatric surgery.

## Methods

### Ethics Statement

All animal experiments in this study were conducted in strict accordance with relevant ethical guidelines and regulations. All procedures were conducted with prior approval from the Danish Animal Experiments Inspectorate, as per the guidelines recommended by the Danish Animal Experiments Inspectorate, approval numbers 2021-15-0201-00831 (University of Copenhagen) and 2023-15-0201-01572 (Aarhus University). At the end of the experiments, all mice were euthanised humanely by cervical dislocation. The study has been reported in accordance with the ARRIVE guidelines [16].

### Animals and Experimental Design

Male C57BL/6JOlaHsd mice (4 weeks old) were purchased from Envigo (the Netherlands) and raised in cages in a specific pathogen-free environment with a diurnal cycle of 12 hours and appropriate temperature (21°C-25°C) and humidity (40%-70%). At 6 weeks of age, mice with similar body weight were randomly divided into the following four groups: low-fat diet group (LFD) fed a low-fat diet; high-fat diet (HFD) group: fed a high-fat diet; LFD + Infection group (LFDinf), fed a low-fat diet and infected with ∼200 3^rd^ stage (L3 stage) *H. p. bakeri* larvae; HFD + Infection group (HFDinf), fed a HFD and infected with ∼200 L3 stage *H. p. bakeri* larvae. Diets were obtained from ssniff Spezialdiäten GmbH and stored at 4°C.

### Feeding and infection with *H. p. bakeri*

In the experimental design, 6-week-old male C57BL/6JOlaHsd mice were fed with either low fat diet (#D12450B, DIO –– 10 kJ% fat, 33% sucrose containing 10 kJ% fat, 20 kJ% protein, 70 kJ% carbohydrates) or high fat diet (#D12492, DIO –– 60 kJ% fat, Lard containing 60 kJ% fat, 20 kJ% protein, 20 kJ% carbohydrates) for 13 weeks. A single dose of 200 3^rd^ stage *H. p. bakeri* larvae was administered by oral gavage to LFDinf and HFDinf group mice at 19 weeks of age and continued a low or high fat diet for 3 more weeks until mice were 22 weeks old.

For all experimental animals, body weight was measured weekly. At the end of the experiment, blood was taken from the submandibular vein, and the liver tissue was harvested for further analysis. Liver tissues were either snap-frozen in liquid nitrogen or fixed with 10% formaldehyde.

## Histology

### Hematoxylin & Eosin staining

Formalin-fixed liver tissues were paraffin-embedded and cut into 5 μm-thick tissue sections. Hematoxylin and eosin staining was performed as per established protocols [17]. Slides were scanned using the VS200 Slidescanner (Evident Scientific, Olympus) scanner at 40× magnification. The histopathological results were evaluated by a certified pathologist specialising in liver pathology at the Odense University Hospital (OUH, Denmark).

### Immunohistochemistry

Formalin-fixed paraffin-embedded (FFPE) liver blocks were sectioned at 3.5 µm thickness and deparaffinised in a xylene ethanol gradient. Slides were incubated overnight in TE buffer (pH 9.0) at 60°C, followed by washing in Tris-buffered saline (TBS) and blocking with 5% casein/TBS for 20 mins. Primary antibody incubation was performed for 2 hours using a rabbit anti-perilipin 2 antibody (1:200, ab52356, Abcam). Slides were then washed in TBS/Tween and the primary antibody was labeled with alkaline phosphatase-conjugated anti-rabbit IgG polymers (BrightVision, Duiven, Holland) for 30 mins, then washed again in TBS/Tween and demineralised water. Signal was visualised with Liquid Permanent Red (DAKO, Glostrup, Denmark). Slides were counterstained with Mayeŕs Hematoxylin, mounted with Aquatex mounting medium (Merck Millipore, Darmstadt, Germany) and scanned at 20× in a VS200 Slidescanner (Evident Scientific, Olympus). For negative controls, a similar protocol was followed, omitting the primary antibody. Digitalised scans were analysed using artificial intelligence-assisted histology through HALO (v4, Indicalabs) and Image J.

### Proteomics

#### Sample preparation – tandem mass tag (TMT) labelling and peptide fractionation

Mouse liver tissue was homogenised with mortar and pestle maintained cold using a mixture of dry ice and 96% ethanol. Approximately 200 µL of 80% methanol in water (both LC-MS grade, VWR) at −20°C was added to liver tissue lysate samples, and the samples were vortexed for 10 s every minute for 5 mins. The samples were incubated at −20°C for 1 hour to precipitate cells and proteins. Precipitated liver cell samples were resuspended in 1% SDS in 12.5 mM HEPES and hydrolysed with probe ultrasonication (Branson Sonifier 250, Branson Ultrasonics., Danbury, USA) with output control fixed at 3 and 30% duty cycle for two rounds of 5-6 pulses each with 30 s incubation on ice between each round. Lysates were centrifuged at 16,000 g at 4°C for 10 mins and supernatant was collected. Briefly, 50 μg of protein were reduced followed by blocking of cysteines, then alkylated proteins were precipitated with pre-chilled acetone at −20°C overnight. Protein sample pellet was obtained by centrifugation at 8000 g for 10 mins at 4°C, air-dried, and subjected to in-solution trypsin (Sigma Aldrich) digestion as previously described [18].

TMTpro Isobaric Mass Tag (18plex) labelling of peptides was performed according to manufacturer’s instructions (Thermo Scientific). Subsequently, labelled peptide samples were separated by strong cation exchange as described previously [18]L. Peptides were purified using PepClean™ C18 Spin Columns (Pierce) and then evaporated using a speed vac.

#### Liquid Chromatography Tandem Mass Spectrometry (LC-MS/MS) analyses

Samples were analysed by nano liquid chromatography (Easy-nLC 1200, Thermo Scientific)-tandem mass spectrometry (Q-Exactive HF-X Hybrid Quadrupole Orbitrap, Thermo Scientific), essentially as previously described [19]. Peptides were trapped on a pre-column (Acclaim PepMap 100 C18, pore size: 100 Å, particle diameter: 3 µm, inner diameter: 75 µm, length: 2 cm, Thermo Scientific) and separated on a reverse phase analytical column (PepMap RSLC C18, pore size: 100 Å, particle diameter: 2 µm, inner diameter 75 µm, length 25 cm, Thermo Scientific) using a 180 mins gradient from 5-90% acetonitrile and 0.1% formic acid at a 270 nL/min flowrate. The MS was operated in positive mode and higher energy collisional dissociation (HCD) with a collision energy (NCE) of 35 was applied for peptide fragmentation. Full scan (MS1) resolution was 60,000 and Automatic Gain Control (AGC) target set at 3×10^6^ with scan range between 392-1,500 m/z. Data-dependent analysis (DDA) was applied to fragment up to 12 of the most intense MS1 peaks. Resolution for fragment scans (MS2) was set at 45,000 with first fixed mass at 110 m/z, maximum injection time 100 ms and AGC target at 1×10^5^. Dynamic exclusion was set at 15 seconds and unassigned and +1 charge states were excluded.

#### Proteomic Database Searches

Database searches were conducted in Proteome Discoverer 3.0 (Thermo Scientific), using the Sequest algorithm on all raw files, merged against *Mus musculus* database containing 16,996 entries retrieved from Uniprot. A maximum of 2 missed cleavages was allowed using trypsin as the enzyme. Precursor and fragment mass tolerance were set at 10 ppm and 20 mmu, respectively. Oxidation of methionine was set as a dynamic modification and static modifications included carbamidomethylation of cysteines and TMT-labels on lysine and the peptide N-terminus. Co-isolation threshold set at 30 %. The identification false discovery rate was set to 0.001. Only proteins with unique peptides were included, and quantitation was based solely on those unique peptides.

### Deworming experimental model

Diet-induced obese (DIO) or DIO control male C57BL/6NTac mice (22 weeks old) were purchased from Taconic (USA) and raised in cages in a specific pathogen-free environment with a 12-hour diurnal cycle, appropriate temperature (21°C-25°C), and humidity (40%-70%). DIO mice with similar body weight were randomly divided into the following three groups: HFD group (high-fat diet as described earlier): HFDinf group (HFD and infection routine as described earlier), and HFDinf + dewormed group (HFDdeworm). For HFDdeworm group, HFD mice infected with ∼200 L3 stage *H. p. bakeri* larvae for 3 weeks were dewormed with oral administration of 100 mg/kg body weight pyrantel pamoate, 2 doses on consecutive days, as previously described. After deworming, mice were observed for an additional 5 weeks while being fed on HFD until they were 30 weeks old. Body weight was measured weekly and at the end of the experiment, blood was collected from the submandibular vein, and the liver tissue was harvested for further analysis. Liver tissues were either snap-frozen in liquid nitrogen or fixed with 10% formalin.

### Interleukin-4 (IL-4)-treated animal model

Six-to-eight weeks old C57BL/6J wild type mice (n=4 per group) were housed in a temperature-controlled room (22°C) with a 12-h light/dark cycle and 55% relative humidity. Food and water were available ad libitum. All research was conducted under the Animals (Scientific Procedures) Act 1986 Amendment Regulations 2012, following ethical review by the University of Cambridge Animal Welfare, under pathogen-free conditions and in accordance with UK Home Office guidelines. *In vivo* IL-4 administration was performed as previously described [20]. Briefly, animal-free recombinant murine IL-4 (AF-214-14, PeproTech) was prepared at 500 μg/mL and complexed with anti-mouse IL-4 antibody (BioXcell, clone 11B11) at a molar ratio of 2:1 (weight ratio 1:5) for 1–2 min at RT. The resulting IL-4/antibody complex (IL-4c) was diluted in PBS to 25 μg/mL IL-4 and 125 μg/mL 11B11. Mice received intraperitoneal injections of 200 μL IL-4c (5 μg IL-4 and 25 μg 11B11) every alternate day and were sacrificed on day 2 (after a single injection) or day 4 (after two injections).

### RT-qPCR

Liver tissue was homogenised in Qiazol lysis reagent using a gentleMACS Dissociator (Miltenyi Biotec, Germany) and filtered through an RNAeasy spin column (Qiagen), including on-column DNAase treatment. RNA was extracted using a commercial miRNAeasy Mini Kit (Qiagen) according to the manufacturer’s guidelines. RNA concentration and purity were measured using a NanoDrop ND-1000 spectrophotometer (NanoDrop Technologies, DE, USA). First-strand cDNA, including gDNA removal, was synthesised using a commercial QuantiTect Reverse Transcription Kit (Qiagen) according to the manufacturer’s instructions and stored at −20°C until further use. RT-qPCR was performed using PerfeCTa SYBR Green Fastmix (Quantabio) on a AriaMx Real-time PCR System (Agilent, USA) under the following conditions: 2 mins at 95°C followed by 40 cycles of 5 s at 95°C and 20 s at 60°C, and finished with 30 s at 95°C, 30 s at 65°C, and 30 s of 95°C again. Primer sequences; Forward: GCCTGCTGAACATCCATGAG; Reverse: TGGTGTCACAGATGGAGTCC. The ΔΔCT method was used to calculate fold changes, relative to the housekeeping gene *Gapdh*. Samples that failed to amplify (CT ≥ 35 in the housekeeping gene) were excluded from further analysis.

### Metabolomics

#### Sample preparation

Mouse liver tissue was homogenised with mortar and pestle maintained cold using a mixture of dry ice and 96% ethanol. Approximately 200 µL of 80% methanol in water (both LC-MS grade, VWR) at −20°C was added to liver tissue lysate samples, and the samples were vortexed for 10 s every minute for 5 minutes. The samples were incubated at −20°C for 1 hour to precipitate cells and proteins. Subsequently, samples were centrifuged 15,000 g for 20 mins at 5°C and the supernatants containing metabolites were collected. The supernatants were vacuum-centrifuged until dry and stored at −20°C.

#### LC-MS/MS-based metabolomics analyses

The samples were analysed by liquid chromatography-mass spectrometry (LC-MS) as previously described [21]. Briefly, the samples were analysed in 20 min gradients on a Vanquish Horizon LC coupled with a Q Exactive Plus Orbitrap MS, both from Thermo Fisher Scientific. The MS analyses were performed in positive MS mode, at an electrospray voltage of 3500 V, and the transfer tube was kept at 300°C. The MS scanned from 70 to 1050 *m/z* in full scan mode. Data-dependent acquisition (DDA) was applied for compound identification, with MS/MS fragmentation of the top 10 peaks, dynamic exclusion of 10 s, AGC of 1 x 10^6^ and an IT of 50 ms. These DDA settings were applied to injections distributed throughout the study, which were used for compound identifications. All other LC-MS analyses were performed by full-scan MS only. The fragmentation was performed stepwise with higher collision dissociation (HCD) at normalised collision energy levels of 20, 40 and 60 eV. Fragmentation scans (MS/MS) were performed at a mass resolution of 17,500 and the full scan MS was operated at high resolution (70,000) and accurate mass (<5 parts per million) to ensure high analytical selectivity.

#### Metabolomics Database Searches

Compound Discoverer 3.3 (Thermo Scientific) was used for identification and calculating the relative abundances of the small molecules in the samples. The identification nodes were predicted composition, the database mzCloud (endogenous metabolites), and Chemspider (Human Metabolome Database, KEGG). The maximum allowed mass deviation between experimental MS data and database values was five parts per million.

### Patient study population

The clinical material analysed in the present study is a part of a larger study conducted in Odense and Esbjerg, Denmark, evaluating the effects of bariatric surgery registered at ClinicalTrials.gov (NCT05291013). In the present sub-study, we analysed plasma samples from age and sex-matched participants with severe obesity without Diabetes (n=15, post-surgery n=12) and healthy lean controls (n=15). The lean controls were requested to be between 30-65 years old, have a normal BMI of 18.5–25 kg/m², to be drug-naïve, have no family history of Diabetes, and exhibit normal levels of HbA1c, fasting plasma glucose and 2-hour plasma glucose following a 75 g oral glucose tolerance test (OGTT). The participants with severe obesity, were requested to be between 30-65 years old, have a BMI of 35–55 kg/m², HbA1c and fasting plasma glucose, and have no family history of Diabetes. Furthermore, they had to meet the Danish eligibility criteria for bariatric surgery (BMI >40 kg/m² or BMI >35 kg/m² with at least one obesity-related comorbidity) and achieve 8% weight loss before referral for bariatric surgery. The patients with severe obesity were permitted to use up to two antihypertensive drugs and one lipid-lowering agent. All participants had normal blood-screening tests of hematologic, renal, and hepatic function and an electrocardiogram (ECG). Prior to involvement, informed consent was obtained from all participants. The protocol was approved by the Regional Committees on Health Research Ethics for Southern Denmark (S-20190162) and complied with the Helsinki Declaration.

#### Study design

At baseline, both groups were examined at two visits. The first visit consisted of anthropometric measurements (height, weight, waist and hip circumference), screening blood sampling, blood pressure, ECG, and a 75 g OGTT. The second visit, which was scheduled at least 2 days after the first, comprised a 4-hour hyperinsulinemic-euglycemic clamp with an insulin infusion rate of 40 mU/m^2^/min^1^ to estimate the M-value as a measure of insulin sensitivity. The plasma samples used in the present study were drawn before initiating insulin infusion. The participants with severe obesity were invited for similar visits 9–12 months after bariatric surgery. Eight females and four males were examined at the follow-up visits. All participants included in this study had improved circulating triglyceride levels 9 months post-bariatric surgery. All visits were performed after an overnight fast, and before the clamp visits, the participants abstained from alcohol, caffeine, and vigorous exercise for 48 hours. Any medication was discontinued one week prior to clamp visits.

### TGM2 ELISA

TGM2 levels in plasma samples were analysed using the Human TGM2 ELISA Kit (EH462RB, Thermo Scientific) according to the manufacturer’s instructions. Briefly, 100 μL of each standard and sample was added to the appropriate wells and incubated for 2.5 hours at RT with gentle shaking. After washing, 100 μL of the biotin conjugate (previously prepared) was added to each well and incubated for 1 hour at RT with gentle shaking Following another wash, 100 μL of streptavidin–HRP solution was added to each well and incubated for 45 mins at RT with gentle shaking. Approximately 100 µL of TMB substrate was then added to each well and incubated for 30 mins at RT in the dark with gentle shaking, followed by the addition of 50 μL of stop solution. The absorbance was measured at 450 nm and 550 nm in a spectrophotometer (FLUOstar OPTIMA).

### Bioinformatics

Analyses of enriched proteins were performed using linear statistical models from *Limma* R package. For the enriched protein analysis using a volcano plot, a design matrix was constructed to model group effects, excluding the LFDinf group to focus on specific contrasts. Two primary contrasts were defined: HFD vs. LFD, representing the “disease effect” associated with high-fat diet, and HFDinf vs. HFD, representing the “treatment effect” of *H. p. bakeri* infection. Contrasts were defined using the makeContrasts() function and applied via contrasts.fit(). Empirical Bayes moderation was then applied using eBayes() to improve variance estimation and statistical power. Significantly differentially altered features were identified using topTable(), with statistical significance defined as an adjusted p-value (Benjamini-Hochberg FDR) <0.05 and an absolute log_2_fold change ≥0.5. Statistical significance was assessed using the *ggplot2* package in R.

*H. p. bakeri*–secreted protein (HES) data used in this study were obtained from previously published LC-MS/MS proteomic analyses [22]. The reported protein list was curated, annotated, and used as input for subsequent functional enrichment and network interaction analyses. Amino acid sequence of each *H. p. bakeri*-secreted protein was retrieved from UniProt (https://www.uniprot.org/), then sequence homology search with the Mus musculus proteome was conducted using the BLAST program. The selected corresponding mouse homolog gene represents the top hit for each entry, excluding search results above 90% or below 20% respectively, as arbitrary thresholds. The corresponding mouse homologs of the top HES anti-inflammatory proteins identified from literature were used to generate the interactome protein-protein interaction (PPI) network) with TGM2.

For the predicted host-parasite interaction, the host protein sequences were annotated with InterPro identifiers from UniProtKB entries, and the parasite protein sequences were analysed for conserved domains using InterProScan v5. All proteins have been identified by sets of Uniform InterPro identifiers. A pairwise Jaccard similarity index was computed for host-parasite protein overlap assessment. For any host protein H and parasite protein P, the Jaccard index, J was defined as:

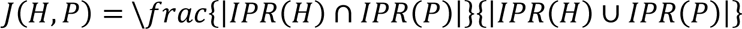

PPIs between Mus musculus (host) and *H. p. bakeri* (parasite, HPOL proteins) were predicted using the PredHPI pipeline developed by KAABIL labs, University of Utah (PredHPI webserver). Searching the HPIDB and STRINGdb repositories under homology thresholds allows for interlog-based prediction. For host and pathogen proteins, a minimum sequence identity of 80%, a minimum coverage of 80%, and an E-value cutoff of 1e-10 were applied to reduce false positives and ensure high-confidence interaction transfer. The predicted host-parasite interaction was visualised in Cytoscape v3.9.

Significantly altered metabolites in the liver tissue of HFDinf mice vs HFD and LFD groups were determined by applying the *Limma* R package. Metabolites were annotated using standardised identifiers (e.g., HMDB, and KEGG Compound IDs) before an overrepresentation analysis based on hypergeometric distribution was performed. Overrepresentation analysis of enriched metabolites was done using the MetaboAnalyst platform (version 6.0).

Protein–metabolite correlation pairs were visualised using the *ComplexHeatmap* R package and correlated using the cor() function in R database and this approach was chosen to quantify linear associations between proteins and metabolites across samples. Metabolite class annotations were derived from the *RefMet* R database, allowing classification into main class and super class categories including lipids, amino acids and organic acids, and nucleic acids. Visualisation of differentially enriched proteins or metabolites was performed using the *pheatmap* R package.

### Statistical analysis

Descriptive statistics are reported as mean ± standard deviations. Unpaired Student T-tests were used for comparisons between the two groups. Multiple group comparisons were performed with a one-way ANOVA. Mann-Whitney U tests (for two groups) or Kruskal–Wallis H tests (for multiple groups) were employed for datasets that did not meet assumptions of normality or homogeneity of variance. P-values were adjusted for multiple testing using the Benjamini-Hochberg method. Pearson’s correlation coefficient was used to evaluate the correlation between groups. P-values <0.05 were considered significant. All data were processed and visualised in R program (version 4.4.2) or GraphPad Prism software, unless stated otherwise.

## Results

### *H. p. bakeri* infection results in reduced lipid loading in the hepatocytes of DIO mice

We generated a DIO mouse model by feeding 4-week-old C57BL/6JOlaHsd wild-type mice a 60% HFD for 16 weeks. They reached a maximum body weight of 54 g after 16 weeks on HFD, representing a significant weight gain of ∼20 g compared with LFD control mice. Infection in DIO mice (HFDinf) or LFD fed mice (LFDinf) was induced by intragastric inoculation of 200 infective L3 stage *H. p. bakeri* larvae, which resulted in an average of 10% weight loss in DIO mice (Fig 1A & Fig S1). This suppression of weight gain is well recognised and has been attributed to the ability of helminth infection to induce a type 2 immune response controlling host inflammatory responses through excretory/secretory products [6, 23, 24] that contain anti-inflammatory molecular determinants (Fig S2). Crucially, infection-induced weight loss is accompanied by reduced hepatic fat accumulation [10]. Consistent with this, we observed a significant reduction in lipid loading in the hepatocytes of HFDinf mice. However, incidentally, the reduced lipid loading was associated only with a decrease in lipid droplet size, not with the number of lipid droplets (Fig 1B). To substantiate further the change in lipid-laden hepatocytes in the liver of HFDinf mice, we stained liver tissues for perilipin-2, a constitutively associated cytoplasmic lipid droplet coat protein [25]. As expected, intense staining was observed at different regions across the liver tissue section of HFD mice, which was reduced significantly in HFDinf mice, whereas LFD control only had sparse accumulation of perilipin staining (Fig 1C).

**Figure 1.**
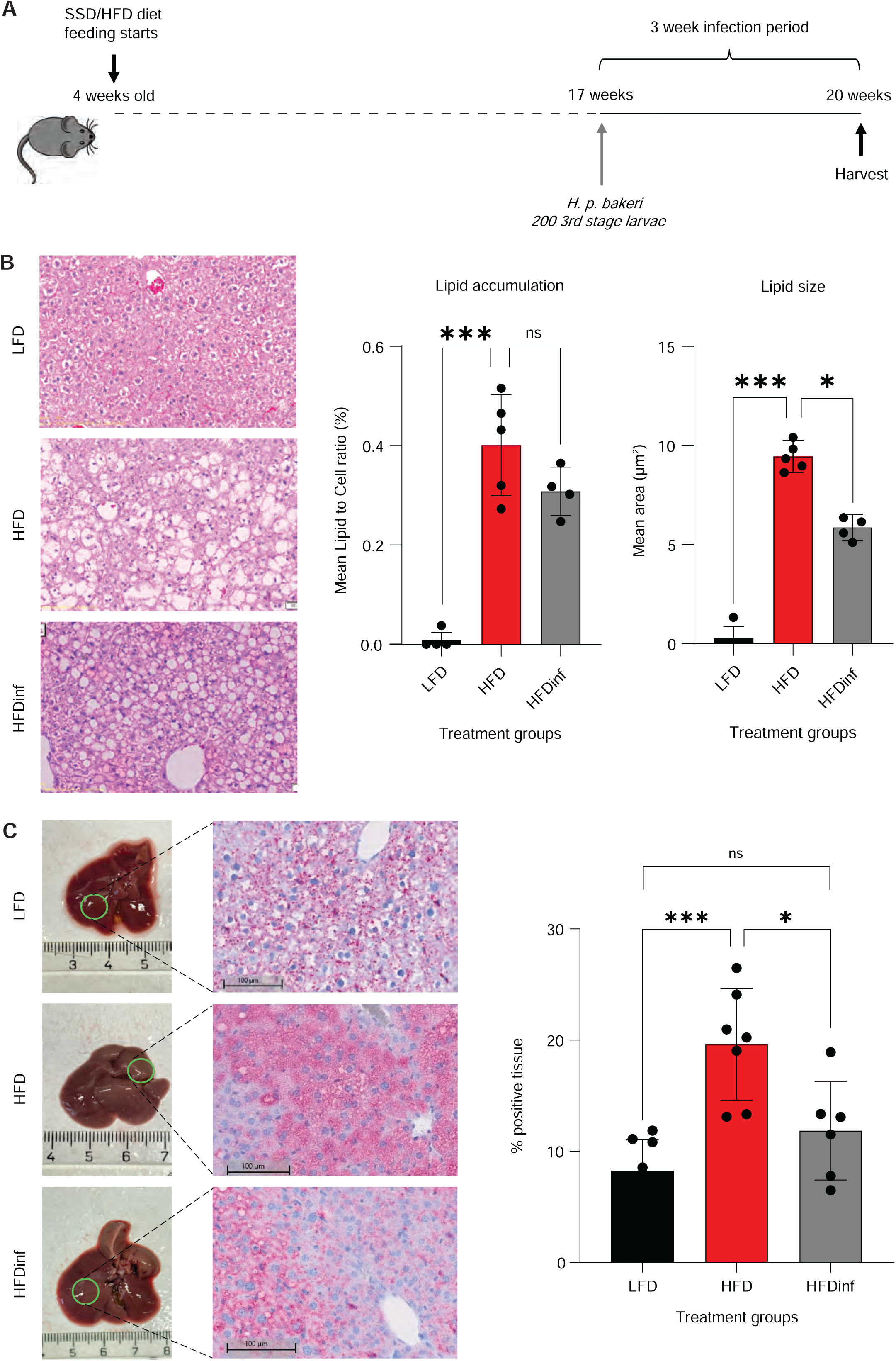
*H. p. bakeri* infection reduce lipid loading in liver hepatocytes of DIO mice. (A) Schematic representation of the timeline of high-fat diet (HFD)-induced obesity in C547BL/6JOlaHsd wild type mice and infection by *H. p. bakeri* (HFDinf). (B) H&E staining displays the relative abundance of lipid accumulation in the liver tissue of HFDinf, HFD, and low-fat diet (LFD) mice. Each dot in the quantification graphs represents an individual mouse per group for all conditions (n=5, LFD and HFD groups) except HFDinf group (n=4). (C) Immunohistochemical analysis showing the intensity of lipid coating perilipin-2 protein staining in the liver tissue of HFDinf, HFD, and LFD mice. Each dot in the quantification graph represents an individual mouse per group for all conditions (n=7, LFD and HFD groups) except HFDinf group (n=6). (B&C) From each scanned slide, six or more different rectangular regions of interest were randomly selected to ensure representative sampling of the tissue. T-test represents difference between the groups. *P < 0.05; ***P < 0.001; ns, not significant.

Collectively, these results validate previous findings on infection-induced changes in hepatic lipid load and provide unique observations that these changes are mostly associated with lipid droplet size.

### Transglutaminase 2 is a novel indicator of infection-mediated regulation of hepatic steatosis in DIO mice

To investigate the mechanistic basis of infection-induced reduction in hepatocyte lipid accumulation, we performed untargeted LC-MS-based proteomic analysis of liver tissue obtained from HFDinf mice versus HFD and LFD controls using TMT labeling + high-resolution mass spectrometry. We identified more than 3800 proteins, and the similarity of protein measurements in LFD, LFDinf, HFD or HFDinf groups was assessed using an unsupervised agglomerative hierarchical clustering (AHC) that grouped replicate values in proximity, indicating negligible variation within treatments (Fig 2A). The LFDinf group, used as a control for diet and infection, was not included in further analysis. Principal component analysis and visualisation of the identified proteins showed a distinctive proteomic status for HFDinf compared to control HFD and LFD mice groups (Fig 2B). Furthermore, deconvolution of the identified protein clusters from HFDinf, HFD, and LFD mice liver samples revealed enrichment patterns in HFDinf that differed from those in the HFD and LFD groups, strongly indicating the creation of a new infection-induced proteomic profile distinct from both HFD- and LFD-induced alterations (Fig 2C).

**Figure 2.**
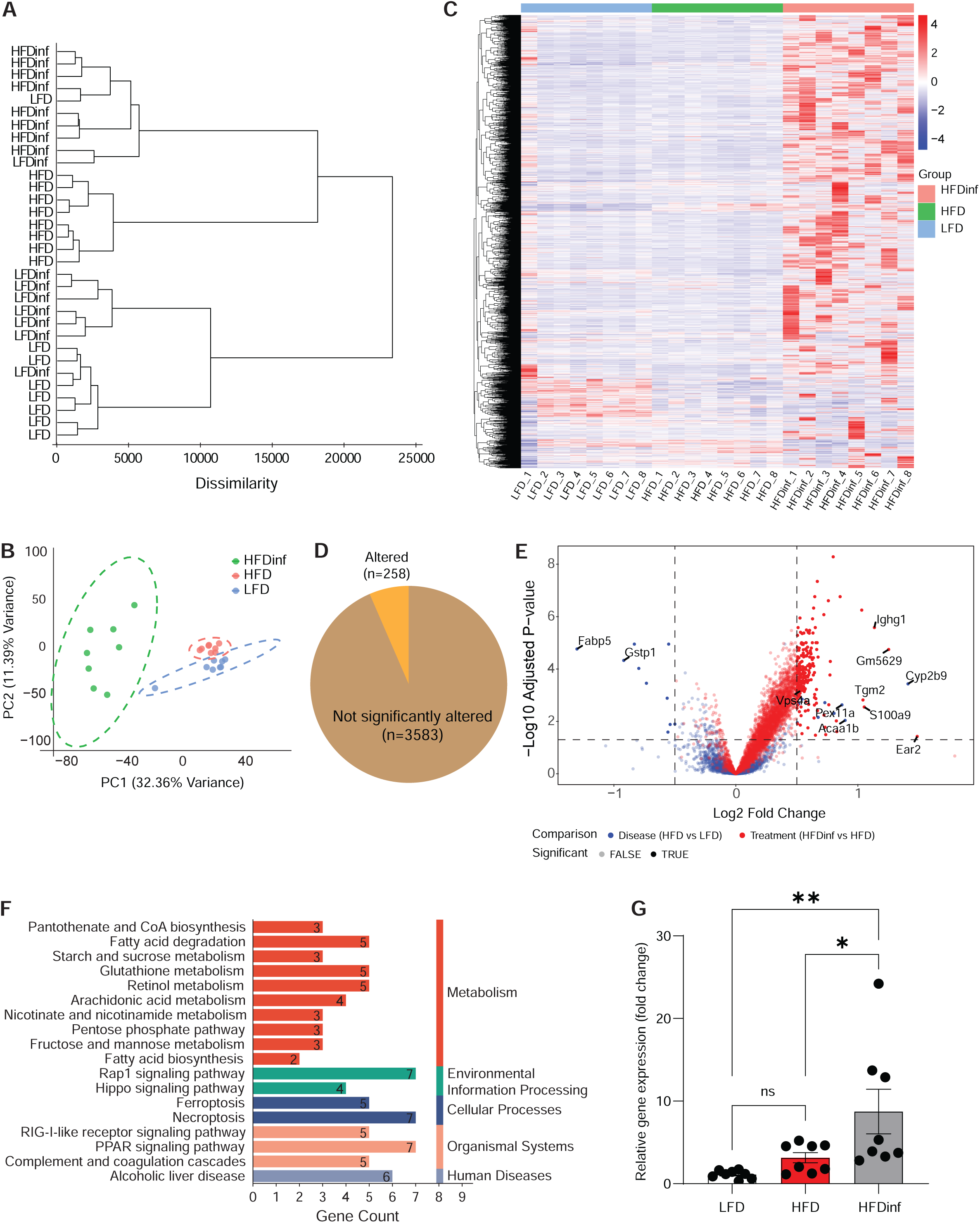
Tandem mass tag (TMT) targeted proteomic analysis of liver tissue from *H. p. bakeri* infection (HFDinf), high-fat diet (HFD), and low-fat diet (LFD) mice. (A) Hierarchical clustering of log_2_-transformed raw protein intensity values identified by TMT targeted proteomics–liquid chromatography–mass spectrometry (LC–MS) on liver tissue from HFDinf, HFD, and LFD mice (n=8 each group). (B) Principal component analysis (PCA) displaying distinct profile of proteins in HFDinf group vs HFD and LFD mice groups. Each dot represents an individual mouse (n=8) per group for different conditions. (C) Differential enrichment of proteins identified from the HFDinf, HFD, and LFD mice groups (n=8 each group) represented by heatmap. (D) Proportion of significantly enriched proteins in the liver tissue of HFDinf mice vs HFD and LFD groups (n=8 each group). (E) Volcano plot showing the top enriched proteins in HFDinf mice vs HFD and LFD groups. (F) Altered pathways of top enriched proteins from proteomic analysis as identified by KEGG analysis. (G) Relative expression of *Tgm2* in liver tissue of HFDinf mice, HFD and LFD mice groups analysed by RT-qPCR. Each dot represents an individual mouse (n=8) per group for different conditions. T-test represents the difference between the groups. *P < 0.05; **P < 0.01; ns, not significant.

Next, we identified 258 proteins to be significantly altered between LFD, HFD, and HFDinf groups from a total of 3841 identities (Fig 2D). All the enriched proteins were subjected to gene ontology (GO) analysis, which most importantly showed a close association of identified proteins with biological processes related to immune regulation (Fig S3A). The top enriched proteins included Immunoglobulin Heavy Constant Gamma 1 (IGHG1), Glycoprotein M6B (GM5629), Cytochrome P450 (CYP2B9), TGM2, Vacuolar Protein Sorting-Associated Protein 4A (VPS4A), S100 Calcium-Binding Protein A9 (S100A9), Peroxisomal Biogenesis Factor 11 Alpha (PEX11A), and Acetyl-CoA Acyltransferase 1B (ACAA1B) (cut-off FC >1.5; p-value adj <0.05, Benjamini-Hochberg correction) (Fig 2E). Intriguingly, IGHG1, GM5629, CYP2B9, TGM2, VPS4A, S100A9, PEX11A, and ACAA1B form an interconnected immune–metabolic network, with TGM2 emerging as a central node linking inflammatory and vesicular signaling pathways to mitochondrial–peroxisomal fatty acid oxidation and lipid homeostasis, thereby influencing hepatic steatosis [26–34]. Further, to determine whether the weight loss in HFDinf mice is accompanied by metabolic regulation, we conducted KEGG analysis, which showed modulation of metabolic pathways by enriched identities of the HFDinf group. A closer look revealed alterations in fatty acid degradation, glutathione metabolism and Peroxisome Proliferator-Activated Receptor (PPAR) signaling pathways, all of which are directly associated with obesity-induced fatty liver disease (Fig 2F). In fact, selected proteins including TGM2 were significantly associated with tissue remodeling, which is a critical feature of MASLD progression (p-value adj <0.05, Benjamini-Hochberg correction) (Fig S3B). This motivated us to validate the differential levels of top enriched proteins in the infection-induced regulation of hepatic steatosis. We performed an RT-qPCR analysis of selected enriched proteins in liver tissues from HFDinf vs HFD and LFD mice groups and ultimately found transglutaminase 2 (*Tgm2*), a negative regulator of adipogenesis [26], to be significantly upregulated in HFDinf versus control HFD and LFD mice groups (Fig 2G).

Taken together, we identified the significance of the novel protein molecule, TGM2, in the *H. p. bakeri*-induced regulation of hepatic steatosis in DIO mice.

### Validation of TGM2 during regulation of hepatic steatosis by *H. p. bakeri* infection using the deworming model

Our results demonstrated that *H. p. bakeri* infection induces upregulation of TGM2 and subsequent reduction in hepatocyte lipid load in DIO mice. To further validate the functional role of TGM2, we hypothesised that removal of *H. p. bakeri* by deworming would lead to downregulation of TGM2 and increased hepatic steatosis. DIO mice at 22 weeks were infected with *H. p. bakeri* (200 L3 stage larvae) and after 3 weeks of infection, treated with pyrantel pamoate for removal of infection or deworming (HFDdeworm). Four weeks after treatment, mouse livers were analysed for changes in TGM2 level and lipid load (Fig 3A). Infection status and removal of infection in HFDdeworm mice were confirmed by examining the presence or absence of eggs in faeces as per our established protocols [35]. Since worm infection and subsequent deworming result in changes in body weight in DIO mice, we monitored their weight changes regularly during infection and treatment with pyrantel pamoate. As expected, HFDinf mice lost approximately 15–20% body weight at the end of 3 weeks post-infection and upon removal of infection by deworming, HFDdeworm mice lost weight in the first week but gained approximately 15–17% body weight over the next 4 weeks (Fig 3B).

**Figure 3.**
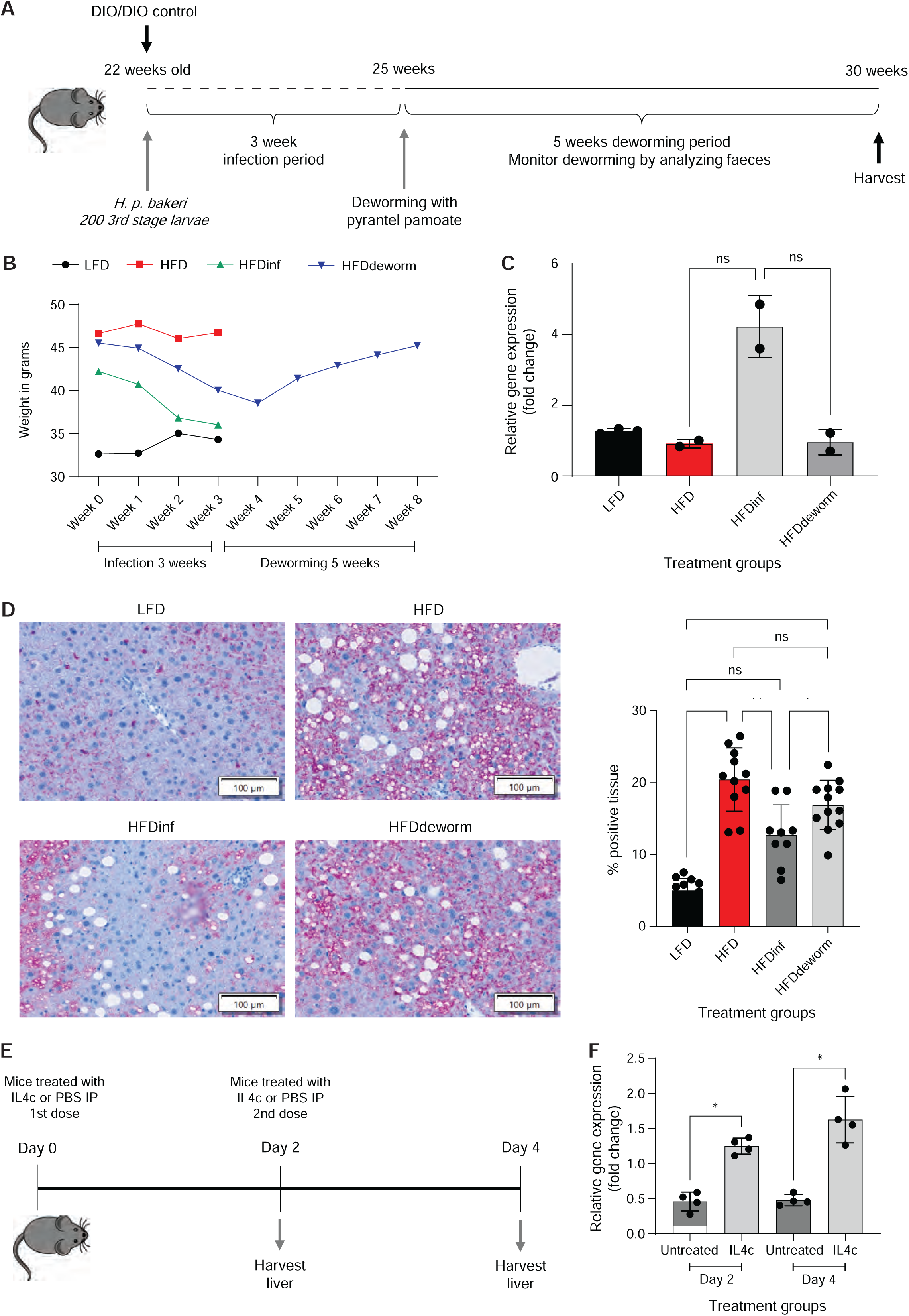
Validation of TGM2 modulation during regulation of hepatic steatosis by *H. p. bakeri* infection using deworming model. (A) Schematic representation showing the timeline of *H. polygyrus* infection in high-fat diet (HFD) mice followed by removal of infection by deworming for a period of 5 weeks. (B) Body weight (grams) changes in low-fat diet (LFD) (n=3), HFD (n=2), *H. p. bakeri* infection in HFD mice (HFDinf) (n=2), and HFDdeworm (n=2) mice groups. (C) Relative expression of *Tgm2* in liver tissue of LFD, HFD, HFDinf, and HFDdeworm mice groups analysed by RT-qPCR. Each dot in the quantification graph represents an individual mouse per group for all conditions (n=2, HFD, HFDinf and HFDdeworm groups) except the LFD group (n=3). (D) Immunohistochemical analysis showing the intensity of lipid coating perilipin-2 protein staining in the liver tissue of LFD (n=3), HFD (n=2), HFDinf (n=2), and HFDdeworm (n=2) mice groups. From each scanned slide, six or more different rectangular regions of interest were randomly selected to ensure representative sampling of the tissue. (E) Schematic representation of mice treated intraperitoneally (IP) with either a single dose of IL4c (interleukin 4-antibody complex) on day 0 or multiple doses on day 0 and day 2, with liver harvested 48h after treatment on day 2 or day 4 for analysis. (F) Relative expression of *Tgm2* in liver tissue of IL4c or PBS control-treated mice analysed by RT-qPCR. Each dot represents an individual mouse (n=4) per group for different conditions. T-test represents the difference between the groups. ****P < 0.0001; **P < 0.01; *P < 0.05; ns, not significant.

Next, to determine if the removal of infection in HFDdeworm mice can revert the TGM2 back to pre-infection levels, we performed a RT-qPCR analysis on the liver tissue from LFD & HFD control vs HFDinf vs HFDdeworm mice. Consistent with the previous observation, infection of HFD mice increased *Tgm2* levels compared with HFD mice, but following deworming, *Tgm2* levels decreased and returned to the pre-infection level (Fig 3C). Further, to correlate changes in body weight and differential *Tgm2* levels with hepatocyte lipid load, we again stained liver tissue from LFD & HFD control, HFDinf and HFDdeworm mice with perilipin-2. As expected, liver tissue staining was more intense in HFD vs LFD mice and upon infection, perilipin-2 staining intensity decreased significantly, indicating an improvement in the hepatic lipid profile in HFDinf mice, accompanied with weight loss and higher *Tgm2* levels as compared to HFD mice. Subsequent deworming resulted in weight gain and an increase in the lipid droplet staining intensity of HFDdeworm mice in line with the change in *Tgm2* levels (Fig 3D).

To further confirm whether alterations in TGM2 are a functional response limited to *H. p. bakeri* infection or part of a broader helminth-induced IL4-mediated type 2 immune response, we analysed changes in TGM2 level in IL4c-treated vs untreated mice. Treatment with IL-4c induced a potent Th2-type immune response in the absence of infection. IL-4c administration led to increased hepatic expression of the macrophage marker F4/80, indicative of macrophage accumulation, along with elevated expression of alternatively activated macrophage markers (Retnla, Mgl2, and Mrc1). Significantly, IL-4c treatment was associated with a robust induction of *Tgm2* expression, suggesting that enhanced *Tgm2* expression is a hallmark of the Th2 immune response rather than a phenomenon specific to helminth infection (Fig 3 E&F and Fig S4).

Altogether, these findings demonstrate that TGM2 is a reliable indicator of hepatic steatosis alterations by helminth infection in DIO mice.

### Direct interaction of *H. p. bakeri* infection-derived secretome and enriched HFDinf mice host proteins

Given the strong evidence linking regulation of TGM2 levels by *H. p. bakeri* infection in hepatic steatosis, we next explored the global interaction landscape between infection-enriched host proteins and the *H. p. bakeri*–secretome. We set out by conducting a GO analysis of the *H. p. bakeri*-secreted proteins previously characterised by LC-MS/MS [22]. Notably, the molecular function classification showed a large set of proteins associated with ATPase-coupled transmembrane transporter activity and ATP-dependent catalytic activities, suggesting that parasite-secreted proteins contribute to active energy-dependent transport and metabolic exchange, potentially facilitating host–parasite communication and adaptation within the infected tissue microenvironment. Within biological processes, we observed a significant association of *H. p. bakeri*-secreted proteins with metabolic processes among others, indicating potential roles in host cell modulation and tissue remodeling (Fig 4A).

**Figure 4.**
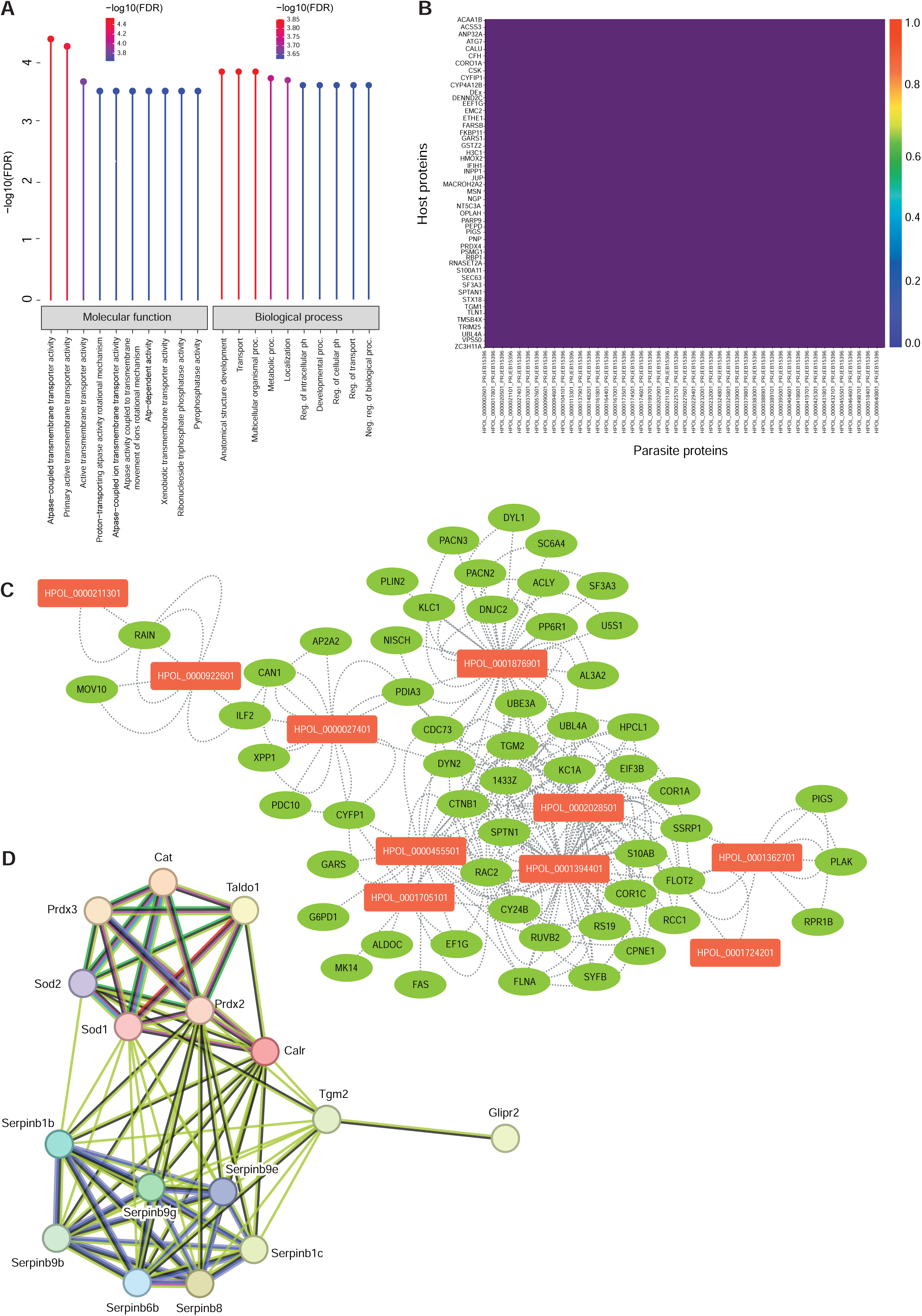
In silico analysis demonstrating the direct interaction between *H. p. bakeri*-derived molecules and infection-enriched proteins in hepatic steatosis. (A) Heatmap showing computed similarity index for host-parasite protein overlap assessment. The similarity scores were assembled in a matrix with host proteins as rows and parasite proteins as columns. Rows and columns of proteins with similar domain architectures were clustered using hierarchical clustering for visualisation of relationships. The heatmap was scaled from 0 (no shared domains) to 1 (complete overlap), and the figure size was adjusted to ensure clear separation and labeling of host proteins along the y-axis. Heatmaps were generated in Python with matplotlib and seaborn libraries. (B) Gene Ontology analysis showing functional classification components, including biological processes and molecular function of *H p. bakeri*-derived proteins, analysed and visualised using PANTHER Classification System. (C) Predicted host-parasite interaction. Each unique protein was represented as a node with an identifier/label, type (host/parasite), and taxonomic annotation. Styled nodes separated the two groups: Host protein images were ovals with green label text, and parasite protein images were hexagons with red label text. Edges were ranked by their confidence score; thicker lines indicate higher confidence predictions. The resulting network was analysed and visualised in Cytoscape v3.9. (D) Interaction of TGM2 with *H. p. bakeri*-secreted molecules with anti-inflammatory properties using STRING 12.0. For protein–protein interaction analysis, *H. p. bakeri* proteins of interest were converted to their corresponding mouse homologs using UniProt to enable cross-species analysis within the STRING Mus musculus database (*H. p. bakeri*, yellow-colored circles and mouse, blue-colored circles). Taldo1, Transaldolase; Psap, Saposin-B-Val; [Prosaposin]; Cat, Catalase; Sod1 & Sod2, Superoxide dismutase [Cu-Zn]; TGM2, Transglutaminase 2; Prdx2 & Prdx3, Peroxiredoxin-2; Calr, Calreticulin; Glpr2, Golgi-associated lipid-binding CAP-domain protein; and Serpins.

The *H. p. bakeri* secretome was then integrated with LC–MS/MS–derived proteomic profile of mouse liver (host) into a unified network analysis pipeline to predict host–parasite molecular interfaces. First, conserved domain architectures were compared using InterPro-based annotations. Pairwise Jaccard similarity indices were calculated for each host–parasite protein pair, yielding a quantitative matrix of shared domain overlap. The resulting heatmap revealed a generally sparse distribution of conserved domains, consistent with limited direct homology between most parasite-derived and host proteins. Nonetheless, discrete clusters of higher similarity were observed, comprising host molecules involved in immune regulation and extracellular signaling, suggesting potential structural convergence or molecular mimicry facilitating host–parasite interaction (Fig 4B).

To further identify potential interspecies molecular interfaces, we performed interolog-based host–parasite protein–protein interaction (PPI) prediction using the PredHPI pipeline. High-confidence interactions were defined using stringent sequence thresholds (≥ 80% identity and coverage; E ≤ 1×10⁻¹⁰) derived from HPIDB and STRINGdb. The resulting Cytoscape-rendered network revealed distinct clusters of interacting *Mus musculus* and *H. p. bakeri* proteins, visualised as confidence-weighted connections between host (ovals) and parasite (hexagons) nodes (Fig 4C). Three *H. p. bakeri* proteins emerged as central hubs including heat shock protein HSP-1 (HPOL_0001876901), actin-4 precursor (HPOL_0002028501), and cytoplasmic actin-2 (HPOL_0001394401), each showing extensive connectivity with host cytoskeletal and chaperone proteins (e.g., FLNA, SPTN1, DNM2). Functional enrichment analysis indicated overrepresentation of GO terms related to protein folding, stress response, and cytoskeletal organisation. HSP-1 carried annotations for ATP-dependent chaperone and stress-adaptive activities, consistent with a role in buffering host proteotoxic stress. In contrast, actin-related proteins were enriched for cytoskeletal remodeling and transcriptional regulation, suggesting parasite-driven modulation of host cell architecture. Beyond their structural and proteostatic roles, these interactions point toward a metabolic regulatory dimension. For example, HSP-1–mediated proteostasis supports mitochondrial and redox homeostasis [36, 37], while actin-dependent cytoskeletal remodeling influences intracellular energy flux through ATP turnover and organelle positioning [38].

Finally, to explore direct links between TGM2 and *H. p. bakeri* immunomodulatory molecules, we performed a STRING v12.0–based network analysis. Secreted *H. p. bakeri* proteins and their mouse homologs with known anti-inflammatory activity were curated through literature mining (Fig S2). TGM2 showed moderate functional connection with the Golgi-associated lipid-binding CAP-domain protein GLIPR2 (the closest mouse homolog of parasite VAL proteins, which is known to modulate host immune response through a variety of ligands including lipids [39]) and superoxide dismutase (SOD2). Weak but significant associations (FDR < 0.05) were also identified with Calreticulin, Thioredoxin Peroxidase (PRDX2), and multiple Serpins (Serpinb1b, Serpinb6b, Serpinb9e, etc.) (Fig 4D).

Collectively, these in silico findings suggest that host TGM2 participates in a conserved immunomodulatory network with *H. p. bakeri*–derived proteins, potentially mediated via GLIPR2- and SOD2-dependent pathways.

### Correlation of *H. p. bakeri* infection-enriched proteins and metabolites to MASLD

After demonstrating differential enrichment of TGM2 by infection in DIO mice with hepatic steatosis, we sought to identify metabolites that may correlate with the *H. p. bakeri* infection-enriched proteins. This is important, since a robust protein-metabolite network and its association with MASLD can further support the role of TGM2 as a predictive biomarker for assessing the *H. p. bakeri* infection-induced regulation of MASLD. To examine this, we performed LC-MS-based metabolomics in DIO mice with altered hepatic steatosis induced by *H. p. bakeri* infection and in control DIO mice. We identified 614 molecular features and after removing uncharacterised detections, around 500 metabolites were selected for further analysis. As with proteomic data, metabolomic measurements from LFD, HFD, and HFDinf group replicates were also found in proximity in an unsupervised AHC (Fig 5A). The identified metabolites in the HFDinf formed a separate cluster compared with the HFD and LFD groups in the partial least squares discriminant analysis, consistent with the proteomic PCA (Fig 5B). Further deconvolution showed variation in metabolite enrichment levels between HFDinf and control HFD and LFD mice liver samples consistent with the pattern observed with the proteomic measurements, which suggest distinct metabolomic changes induced by infection (Fig 5C). Subsequently, we identified 151 metabolites that significantly varied between HFDinf and control HFD and LFD groups out of the total 500 characterised metabolites (cut-off FC >1.5; p-value adj <0.05, Benjamini-Hochberg correction) (Fig 5D). KEGG pathway analysis of the enriched metabolites revealed modulation of metabolic pathways, including nicotinate & nicotinamide, glutathione, vitamin B6 metabolism and others directly associated with MASLD pathology [40–42] (Fig 5E). Next, to delineate the functional relationships between hepatic proteins and metabolic reprogramming in diet-induced obese (DIO) mice infected with *H. p. bakeri*, we performed an integrated protein–metabolite correlation analysis. Pairwise Pearson correlation coefficients (r > 0.7; adjusted *p* < 0.05, Benjamini–Hochberg correction) were computed between significantly altered proteins and metabolites, and the resulting correlations were visualised as a clustered heatmap (Fig 5F). Distinct correlation patterns were observed across major metabolite classes, including amino acids and peptides, fatty esters, pyridine alkaloids, fatty acyls, pyrimidines, purines, glycerolipids, and carbohydrate derivatives. When grouped into higher-order chemical superclasses (lipids, carbohydrates, organic acids, and nucleic acid derivatives), the data revealed two opposing regulatory modules, one enriched in amino acid and peptide-related metabolites that showed positive correlations with infection-induced proteins, and another comprising lipid-derived species that exhibited inverse correlations (Fig S5A). Positive correlations between amino acid–associated metabolites and infection-enriched proteins implicated in redox regulation, mitochondrial maintenance, and cytoskeletal organisation suggest enhanced proteostatic and bioenergetic adaptation under helminth infection. In contrast, the inverse correlations observed for fatty esters and long-chain fatty acyls indicate reduced lipid accumulation and a shift toward increased lipid utilisation in infected livers. Notably, metabolite classes linked to nucleotide metabolism (pyridines, purines, and pyrimidines) displayed infection-specific correlation signatures, suggesting a concurrent modulation of redox balance and biosynthetic flux. Further examination of the protein-metabolite correlation map revealed metabolites involved in TGM2-specific interactions, including L-hexanoylcarnitine, nicotinamide ribotide, alpha-ketoglutaric acid, N-acetyl-L-2-aminoadipic acid, troxipide, and L-valine (Fig 5G & Fig S5B). Importantly, some of these metabolites that strongly interacted with TGM2 (Pearson coefficient r≥0.7; p-value adj <0.05, Benjamini-Hochberg correction) were associated with MASLD. For example, plasma concentrations of N-Hexanoylcarnitine have been found to be higher in MASLD patients [43]. Similarly, circulating levels of alpha-ketoglutaric acid were higher in MASLD patients than in healthy individuals [44].

**Figure 5.**
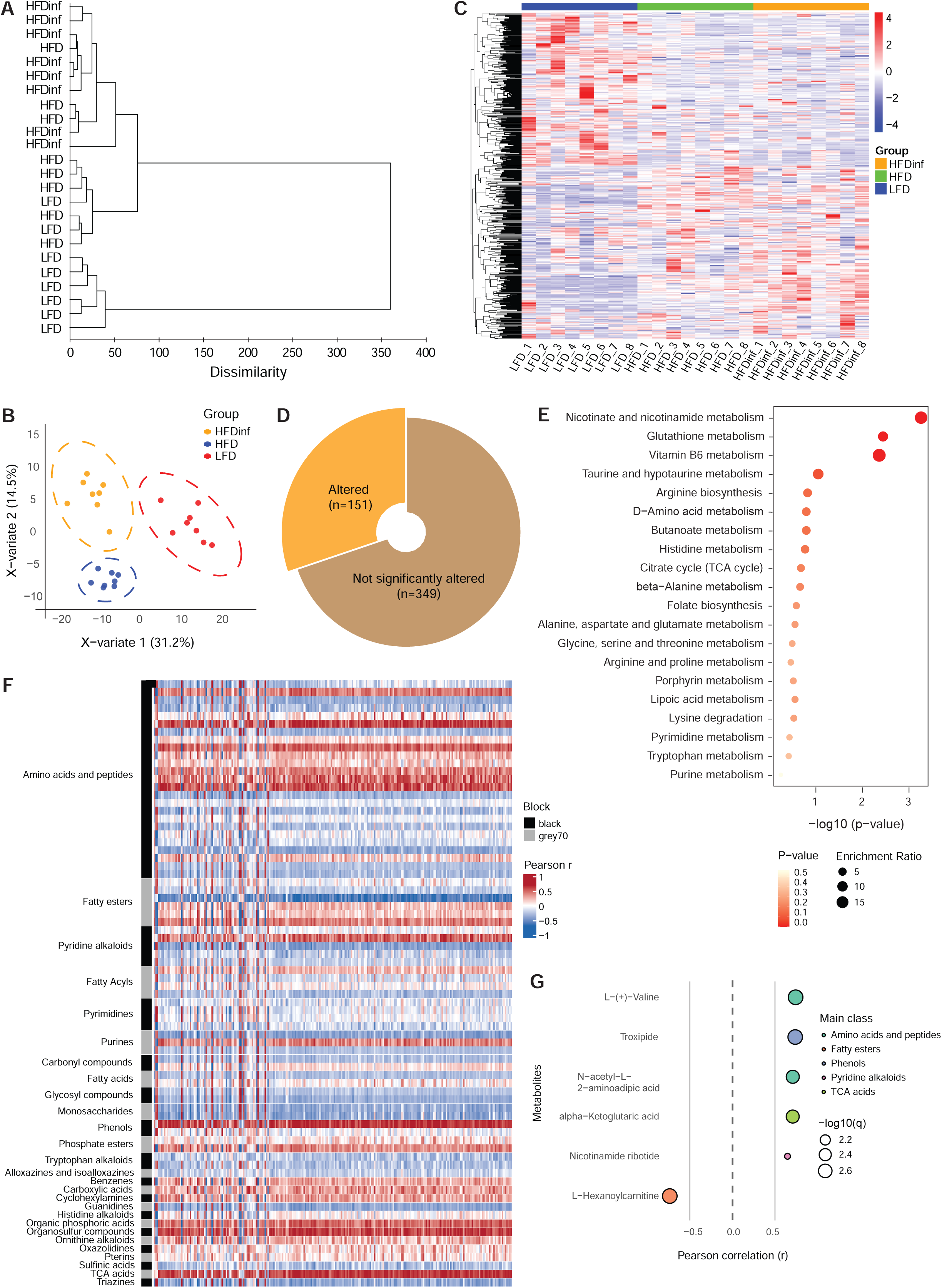
Metabolomic and bioinformatic analysis showing TGM2 as a potential biomarker of MASLD. (A) Hierarchical clustering of log_2_-transformed raw metabolite values identified by metabolomics on liver tissue from *H. polygyrus* infection in high-fat diet (HFDinf), high-fat diet (HFD), and low-fat diet (LFD) mice by liquid chromatography–mass spectrometry (LC–MS). (B) Partial least squares-discriminant score plot analysis (PLS-DA) showing the distinct profile of metabolites in HFDinf group vs the HFD and LFD mice groups. Each dot represents an individual mouse (n=8) per group for different conditions. (C) Differential enrichment of metabolites identified from the HFDinf, HFD, and LFD mice groups (n=8 each group) represented by a heatmap generated using the log_2_-transformed peak intensities already normalised by Z-scoring. (D) Proportion of significantly altered metabolites in the liver tissue of HFDinf mice vs HFD and LFD groups (n=8 each group). (E) Overrepresentation analysis of enriched metabolites revealed metabolic pathways strongly associated with MASLD/MASH progression. (F) Correlation of enriched proteins and metabolites, the correlation matrix was visualised with a heatmap. A pairwise correlation analysis was conducted across the integrated proteomics and metabolomics datasets. Pearson’s correlation coefficients were calculated between all protein–metabolite pairs. Rows represent metabolites and columns represent proteins. (G) Correlation significance of TGM2 with key metabolites. Correlation significance threshold set at |r|>0.5 (adjusted P<0.05), identified significant pairs which were prioritized. All processed data were visualised in R (version 4.4.2).

Together, the results indicate a strong association between top enriched proteins, including TGM2 and metabolites to MASLD and importantly, TGM2 could serve as a marker for monitoring the status of MASLD progression in DIO mice with parasitic worm infection.

### TGM2, a unique predictive marker of worm infection-induced regulation of hepatic steatosis

Given the compelling findings on the differential TGM2 levels in the regulation of hepatic steatosis by *H. p. bakeri* infection and their association to MASLD, we wanted to determine whether TGM2 is a specific response of the worm-induced resolution of hepatic steatosis or a general clinical predictor of improvement in hepatic lipid profile. To study this, we analysed soluble levels of secreted TGM2 protein in obese patients before surgery and 9-12 months after bariatric surgery, which improved hepatic lipid profile (Fig 6A). At baseline, participants with severe obesity had higher weight, BMI, waist circumference and slightly higher HbA1c and fasting plasma glucose, whereas insulin sensitivity (M-value) was lower compared with the healthy lean controls (Table S1). After bariatric surgery, weight, BMI and waist circumference were markedly reduced, whereas HbA1c and fasting plasma glucose improved. In addition, bariatric surgery increased insulin sensitivity (M-value) (Table S1). Further, the improvement in hepatic lipid profile was confirmed by the analysis of circulating triglycerides, a marker of hepatic steatosis, and only matched obese patients who had a decrease in plasma triglyceride levels following bariatric surgery were selected for further analysis (Fig 6B). ELISA measured circulating TGM2 levels in paired plasma samples obtained from individuals with obesity who underwent bariatric surgery (n=12) vs pre-operative obese patients (n=12) and control lean individuals (n=15). Consistent with observations in uninfected mice, the TGM2 levels were not significantly altered in individuals with severe obesity compared with lean individuals. Apparently, TGM2 levels did not change in response to the weight-loss induced by bariatric surgery in the participants with severe obesity (Fig 6C), indicating that the alteration of TGM2 following improvement of hepatic lipid profile is a unique functional response to the regulation of hepatic steatosis by parasitic worm infection.

**Figure 6.**
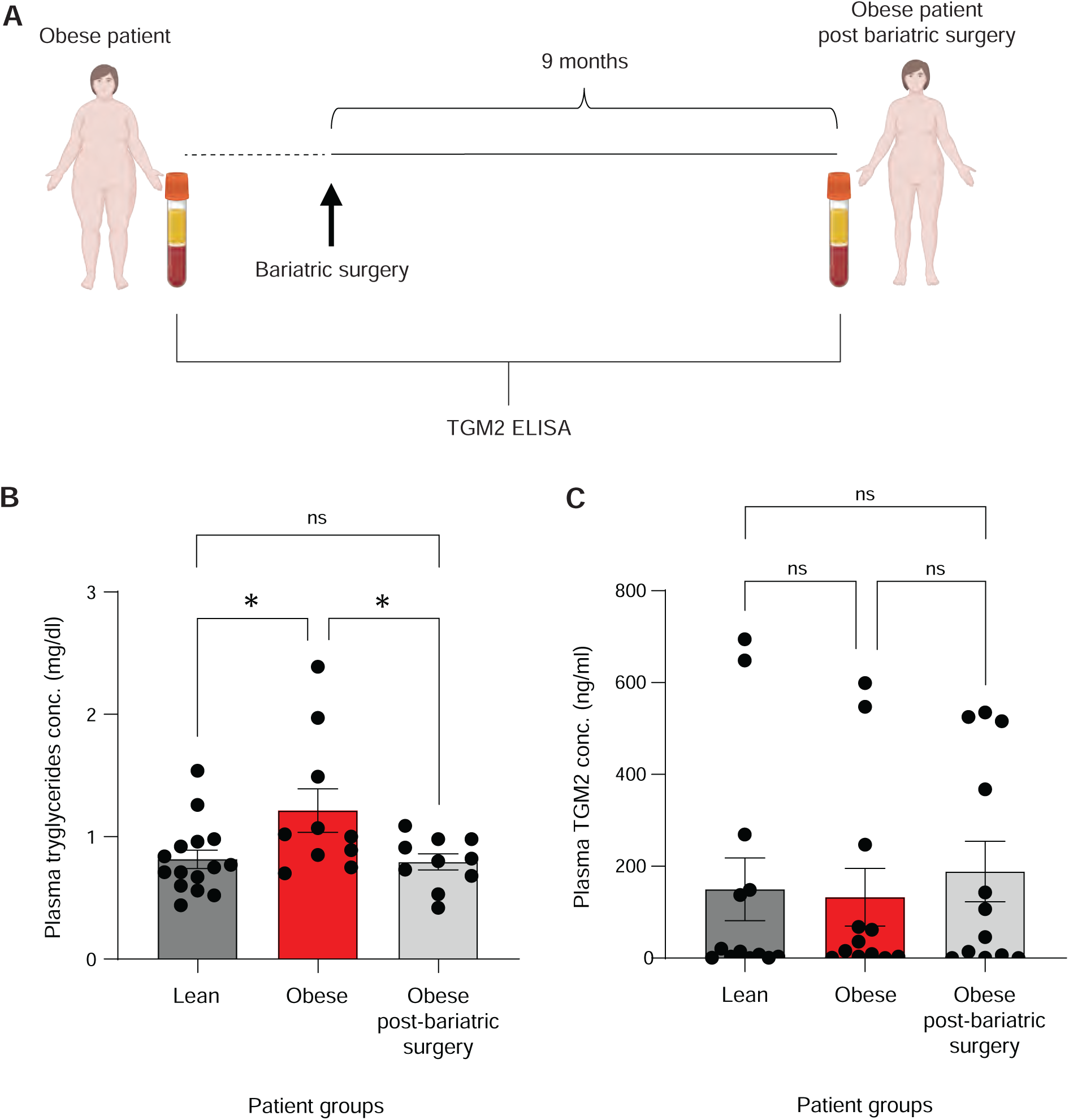
Analysis of TGM2 as a predictive marker of hepatic lipid profile in baseline obese patients and after intervention. (A) Schematic representation of the experimental set-up. Blood plasma was collected from obese patients before surgery and 9 months after bariatric surgery with lean individuals as controls. (B) Triglyceride levels in lean individuals, baseline obese, and obese patients after bariatric surgery. Each dot represents an individual patient’s plasma sample from the obese and obese patients post bariatric surgery group (n=12 each group), except for lean individuals (n=15). (C) Circulating levels of TGM2 in lean individuals, baseline obese, and obese patients after bariatric surgery. Each dot represents an individual patient’s plasma sample from obese and obese patients post bariatric surgery (n=12 each group), except for lean individuals (n=13). T-test represents the difference between the groups. *P < 0.05; ns, not significant. Supplementary figure 2. Identification of top immunomodulatory molecules from *Heligmosomoides polygyrus bakeri* (*H. p. bakeri*) excretory secretory products (HES) by targeted proteomics and in silico analysis, visualized by SankeyMATIC.

## Discussion

Helminth infection resulting in weight loss in DIO mice is accompanied by metabolic homeostasis, which has been implicated on the immunometabolic events triggered by infection-derived products in the host [6, 10, 23, 24]. Although previous studies have described the nature of infection-induced immunometabolic changes in the gut and liver, this is the first-of-its-kind study to provide a detailed account of the proteomic and metabolomic alterations and to identify a molecular target that accurately indicates the regulation of infection-induced metabolic changes in the liver. In this study, we confirmed steatotic changes in the liver and identified TGM2 as a predictive marker of *H. p. bakeri* infection-induced regulation of hepatic steatosis. We validated the differential levels of TGM2 in lipid homeostasis using a deworming and IL4c-treated mice model. Further, we confirmed the TGM2 changes are a unique feature of worm-induced regulation of hepatic steatosis by analysing a patient cohort of lean vs obese vs post bariatric surgery obese patients. Collectively, we provide strong evidence that differential TGM2 levels predict improvement in the hepatic lipid profile of obese infected mice, with potential implications for MASLD pathology. Following *H. p. bakeri* infection, DIO mice lost 15-20% of their body weight, accompanied by a decrease in the size, but not the number, of lipid load, similar to observations in mice and humans following caloric restrictions [45]. In line with the weight change, we expected that *H. p. bakeri* infection in DIO mice would mitigate HFD-induced changes and potentially restore the hepatic proteomic landscape to a baseline state resembling that of the LFD group. In contrast, we observed the development of a distinct proteomic profile in the livers of DIO mice following *H. p. bakeri* infection (Fig 2). Although, the exact mechanistic understanding of this different proteomic status in infected DIO mice remains to be elucidated, it may partly account for the divergence in TGM2 levels observed between HFDinf and LFD control groups. This observation also indicates that alteration in TGM2 levels are a unique function of helminth infection.

Alterations in TGM2 levels in relation to the status of hepatic steatosis were convincingly supported by the deworming experiment, where improvement or worsening of hepatic lipid load following infection and subsequent deworming in DIO mice was consistently reflected by corresponding changes in TGM2 levels (Fig 3). A limitation of this experimental set-up was the small number of mice per treatment (n=2) or control arm (n=3). Nevertheless, given that the deworming was conducted in a different strain of C57BL/6NTac in-bred mice and fed with a HFD different from the previously used experimental setup, this emphasises the reproducible nature of infection-induced changes in TGM2 levels. As HFD-induced obesity leads to gut dysbiosis, resulting in increased permeability of intestinal wall [46–50] and the infection-mediated observations from our study, it is highly likely that *H. p. bakeri*-secreted products may traverse across the intestinal wall, gain access to the portal vein through the small blood vessels, and eventually localise in the liver. Due to the immunomodulatory nature of *H. p. bakeri*-secreted products, they may attenuate hepatic steatosis, as indicated by alterations in the levels of TGM2.

Our *in silico* analysis suggested that *H. p. bakeri* co-opts host metabolic and immune networks through chaperone-mediated stress tolerance and cytoskeletal mimicry, thereby enabling immune evasion and tissue adaptation. Among its secreted molecules, *H. p. bakeri* VAL proteins, such as Hp-VAL4, have been shown to exert critical immunomodulatory functions by binding to a variety of ligands, including sterol and palmitate [39]. Consistent with this, our interaction analysis predicted a moderate association between host TGM2 and GLIPR2 (the closest mouse homolog of *H. p. bakeri*-secreted VAL proteins) (Fig 4), supporting the hypothesis that *H. p. bakeri*-secreted products could influence host TGM2 levels and in turn, hepatic lipid metabolism. The integrated proteomic–metabolomic network further supports this concept showing that helminth infection induces coordinated remodeling of amino acid and lipid metabolism in obese hosts, coupling immune modulation with improved hepatic energy balance and reduced steatosis (Fig 5). Importantly, within this framework, host TGM2 emerges as a potential integrator of these metabolic and immunoregulatory effects. Previous studies have demonstrated that TGM2 regulates hepatic steatosis by controlling autophagy and maintaining mitochondrial homeostasis in obese mice [51]. Together, these findings suggest that *H. p. bakeri* VAL proteins could directly influence hepatic steatosis through TGM2-dependent mechanisms. However, the role of lipid binding in the immunomodulatory functions of VAL proteins is yet to be established and although this hypothesis of host-parasite interaction mediated by TGM2 is plausible, more functional validation studies are required in TGM2 KO animal models to understand if infection-induced regulation of hepatic accumulation of lipid droplets is indeed mediated by TGM2 directly or by alternative pathways.

We compared individuals with obesity before and after bariatric surgery to determine whether differential levels of circulating TGM2 reflect a general functional response to improved hepatic lipid profile or a predictive marker unique to *H. p. bakeri* infection. The main reason for selecting this model was that bariatric surgery has been shown to convincingly reduce liver enzymes, and improve lipid and glycemic profiles in individuals with obesity, emphasising its effectiveness as a treatment strategy for morbid obesity [52]. The absence of a significant change in TGM2 levels between individuals with obesity before and after bariatric surgery (Fig 6) supports the conclusion that changes in TGM2 levels are a unique function of *H. p. bakeri* infection, potentially pointing toward their larger utility in the spectrum of MAFLD/MASH pathology in the subpopulation of patients infected with helminths. In fact, the correlation between top enriched proteins and metabolites and their association with functional pathways of MASLD (Fig 5F&G) is critical knowledge to expedite the development of therapeutic interventions targeting the early stages of MASLD progression. More importantly, the identification of TGM2 and its strong association to MASLD pathways can be expected to provide impetus for prospective investigations on using mimetics of *H. p. bakeri*-derived immunomodulatory molecules or similar molecules as an early intervention next-generation therapeutics for the resolution of MAFLD progression and monitoring of therapy response by measurement of changes in the level of TGM2.

In conclusion, our findings identify TGM2 as a unique predictive marker of improved hepatic lipid profile and provide additional evidence that chronic worm infection protects against obesity-induced hepatic steatosis in mice. Therefore, mimetics of *H. p. bakeri*-derived products, combined with changes in TGM2 levels, may offer novel therapeutic opportunities for preventing the progression of hepatic steatosis to severe forms of liver pathology.

## Acknowledgements

This work was supported by the Lundbeck Foundation (Grants R400-2022-1152 to P.N. and R400-2022-1239 to A.R.W). V.I.C. is grateful to Hørslev Fonden (grant 41301763.1) for supporting this work. G.B., M.D. and A. V-P were supported by the British Heart Foundation (grant number RG/F/23/110110) and the Medical Research Council (grant number MC_UU_00039). We thank Sönke Detlefsen (Dept. of Pathology, University of Southern Denmark) for examining the mice liver tissue H&E staining to confirm hepatic steatosis. We also thank Rick Maizels (University of Glasgow) for granting permission to use published *Heligmosomoides polygyrus bakeri* secretrome proteomics data and for his valuable feedback on the manuscript. We acknowledge excellent technical assistance from Kaja Søndergaard Laursen.

## Author contributions

Conceived, V.I.C.; Conceptualization, V.I.C.; Methodology, S.F., M.S., G.B., M.D.dC., J.P., and V.I.C.; Investigation and formal Analyses, S.F., M.S., R.S., G.B., M.D.dC.,M.D., L.M., K.J.K., V.M., J.P., and V.I.C.; Resources, M.D.dC., C.B.J., A.V-P., K.H., J.P., P.N., A.R.W., and V.I.C.; Writing – Original Draft, V.I.C.; Writing – Review & Editing, S.F., M.S., R.S., G.B., M.D.dC., M.D., L.M., K.J.K., C.B.J., A. V-P., K.H., J.P., P.N., A.R.W., and V.I.C.; Supervision, J.P., P.N., and V.I.C.; Project Administration, V.I.C.; Funding Acquisition, P.N., A.R.W., and V.I.C.

---

**Supplementary figure 1.**
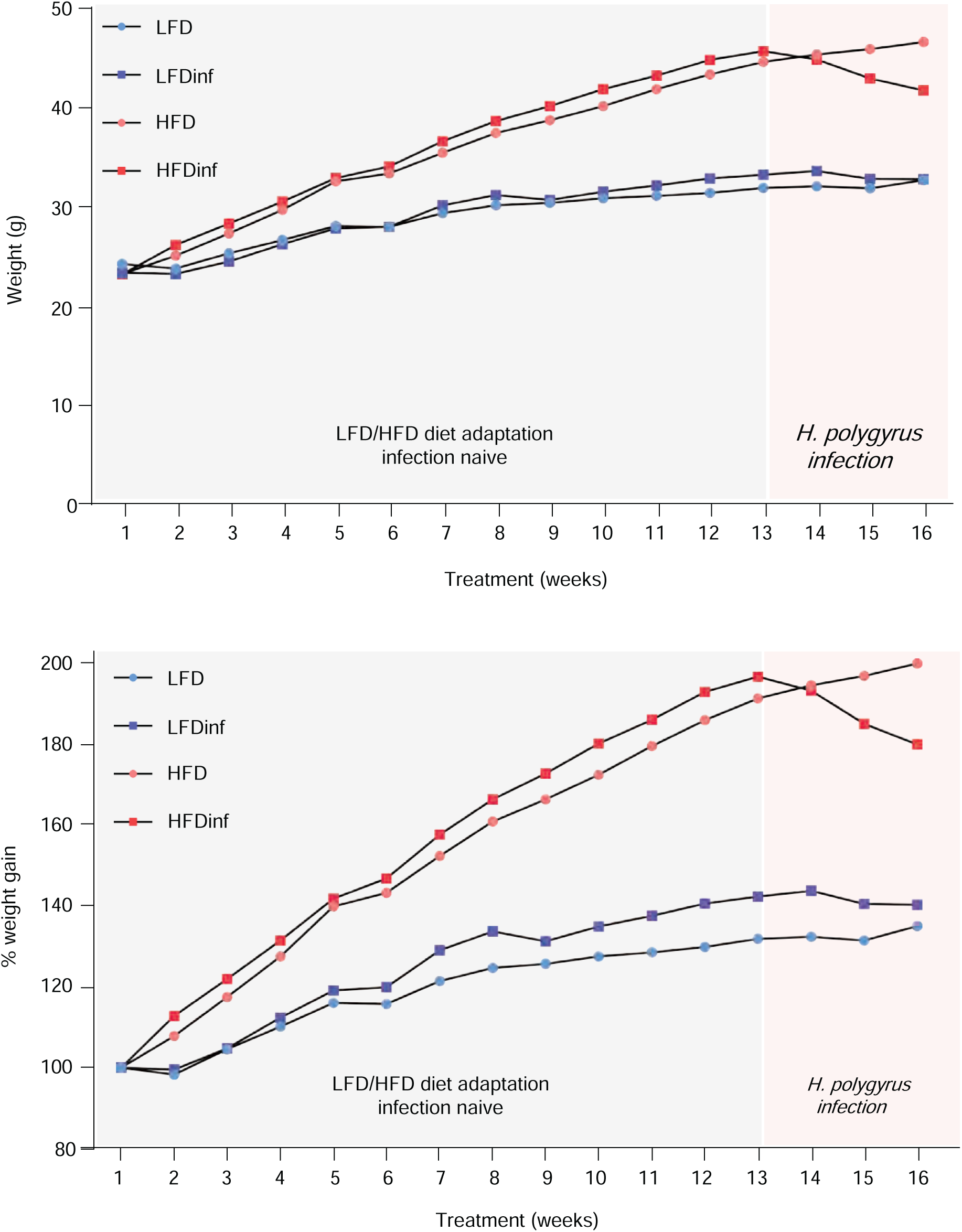
Body weight (grams) changes in C57BL/6 mice that were fed on different diets including low fat diet (LFD), LFD + infection with *Heligmosomoides polygyrus bakeri* (LFDinf), high fat diet (HFD), and HFD + infection with *Heligmosomoides polygyrus bakeri* (HFDinf).

**Supplementary figure 2.**
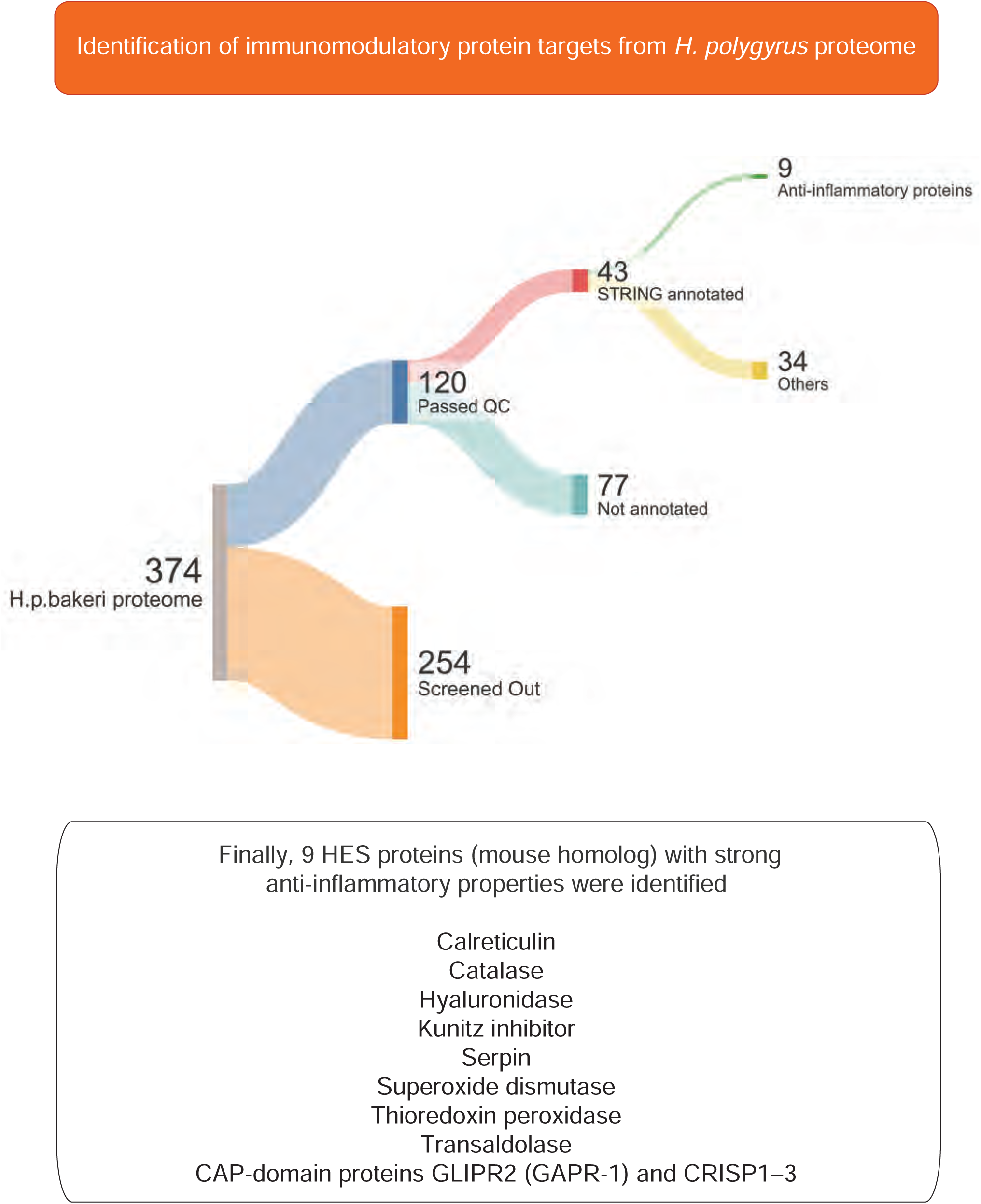
Identification of top immunomodulatory molecules from *Heligmosomoides polygyrus bakeri (H. p. bakeri)* excretory secretory products (HES) by targeted proteomics and in silico analysis, visualized by SankeyMATIC.

**Supplementary figure 3.**
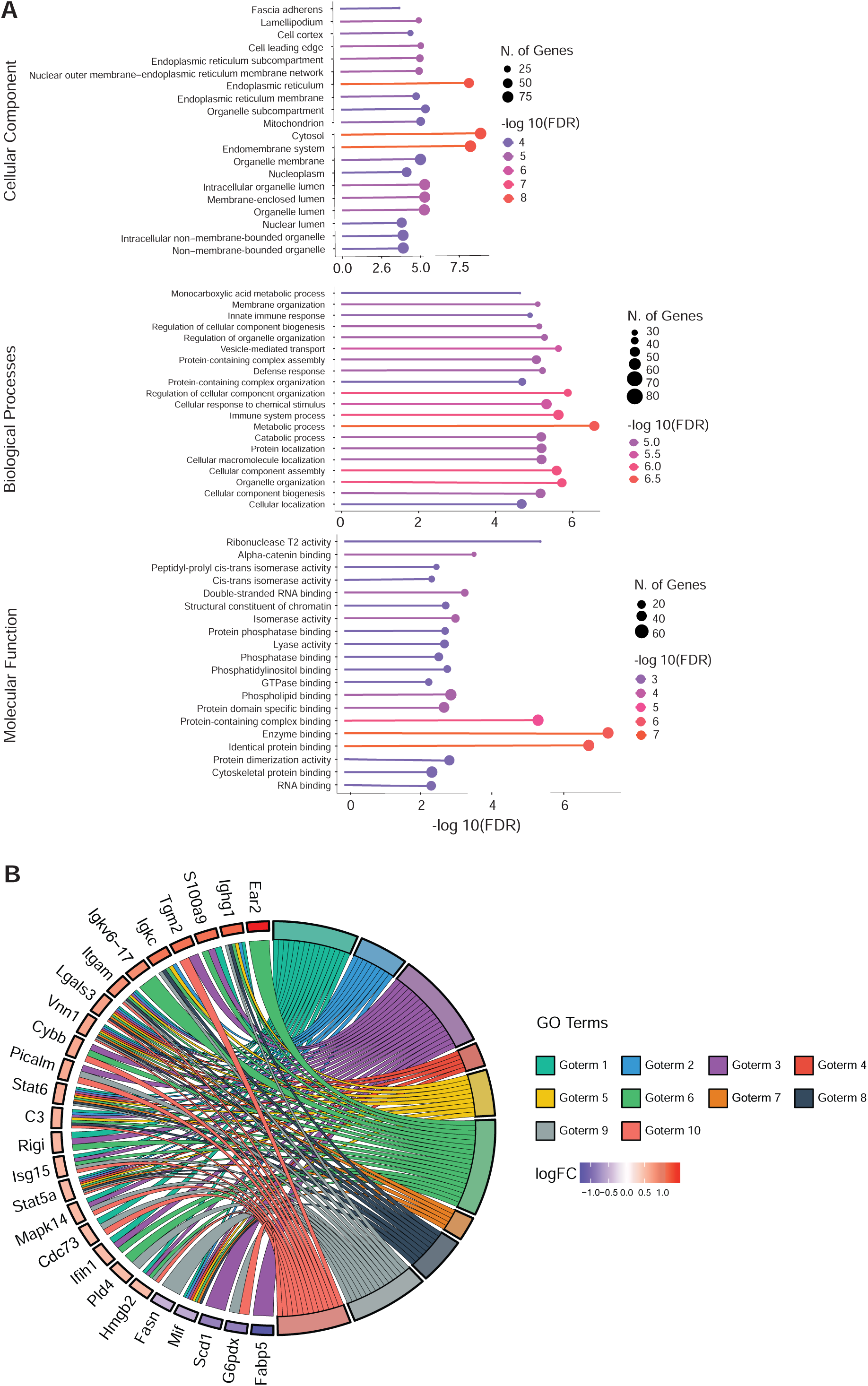
(A). Gene Ontology (GO) analysis showing functional classifications, Cellular Component, Biological Processes, and Molecular Function of top enriched proteins identified by high resolution isoelectric focusing + tandem mass tag mass spectrometry in HFDinf (*Heligmosomoides polygyrus bakeri*-infected diet-induced obese mice) vs high fat diet (HFD) and low-fat diet (LFD) controls. (B). GO-chord network plot showing association of transglutaminase (TGM2) and other enriched proteins to GOTERM biological processes in HFDinf vs HFD & LFD controls. GOTERM 1-10 are biological processes as outlined in panel A.

**Supplementary figure 4.**
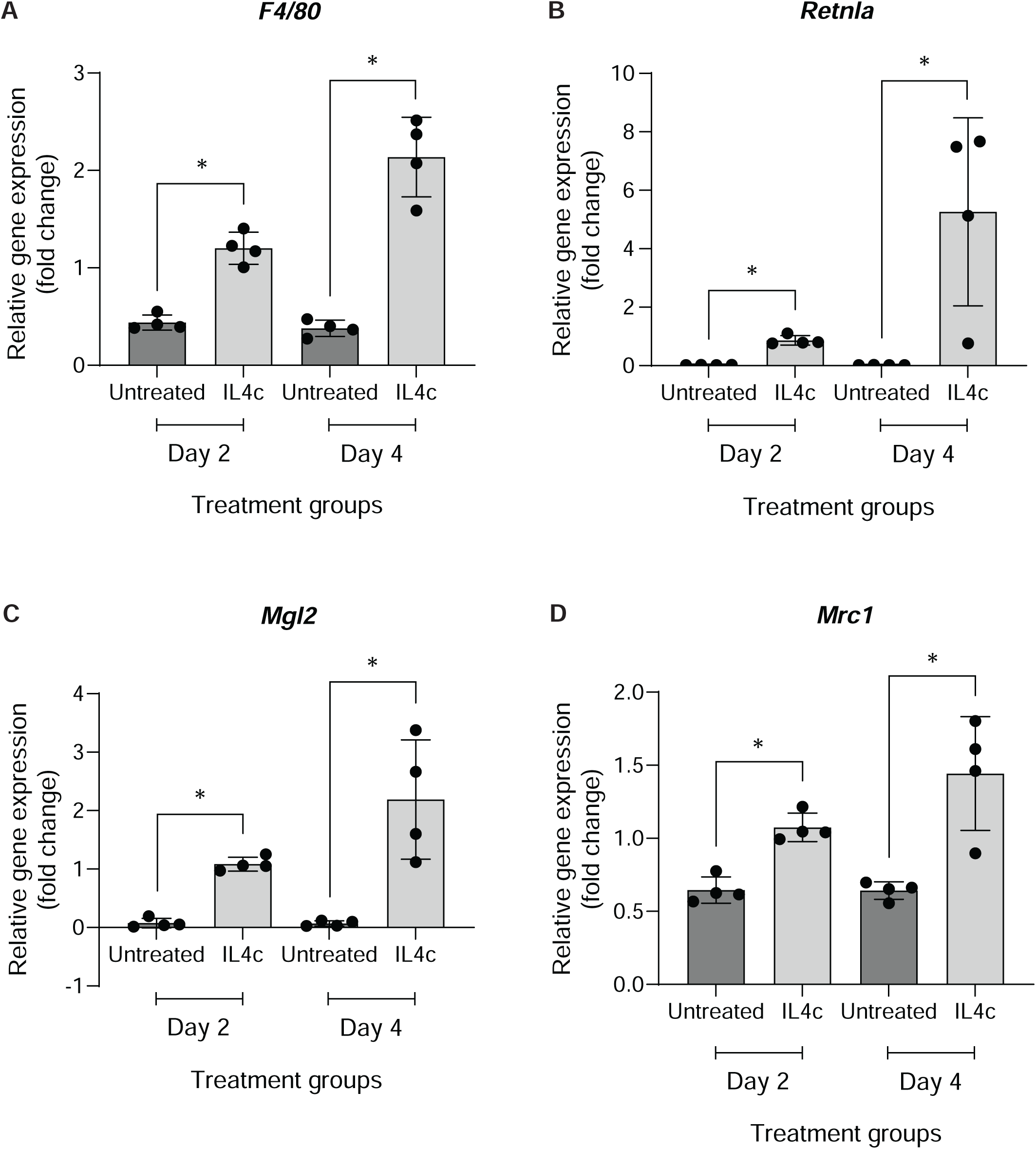
Relative expression of (A) *F4/80*, (B) *Retnla* (Resistin-like molecule alpha), (C) *Mgl2* (macrophage Galactose-type Lectin 2) and (D) *Mrc1* (mannose Receptor C-Type 1) genes in liver tissue of mice treated with IL4c (interleukin 4-antibody complex) or PBS analyzed by RT-qPCR. Each dot represents an individual mouse per group for all conditions (n=4). T-test represents difference between the groups. *P < 0.05.

**Supplementary figure 5.**
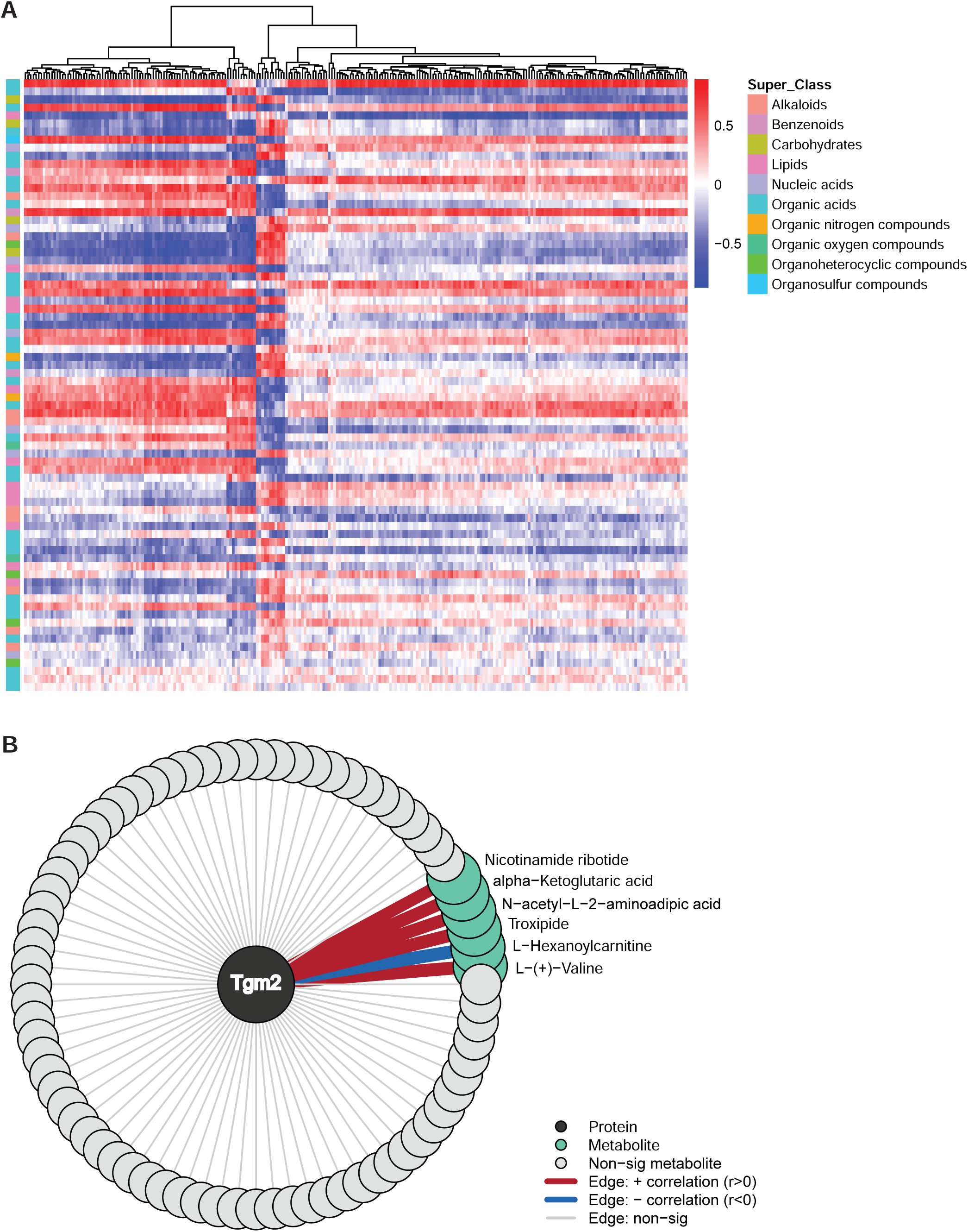
(A). Corrleation of proteins and metabolites significantly enriched in infected diet-induced obese mice (HFDinf) vs high-fat diet (HFD) and low-fat diet (LFD) grouped by RefMet Super Class. (B). Network plot showing the association of 6 top enriched metabolites in HFDinf mice vs HFD and LFD groups to transglutaminase 2 (TGM2).

**Supplementary Table 1.** Table showing patient demography and biochemical parameter measurements.

|  | **Normal** | **Obese baseline** | **Obese post bariatric surgery** | **p-value**<br>**(Obese post-surgery vs baseline)** |
| --- | --- | --- | --- | --- |
| **Participants**<br>**(female/male)** | 15 (9/6) | 12 (8/4) | 12 (8/4) |  |
| **Age** | 46.9 ± 2.5 | 47.3 ± 2.5 | 49.4 ± 2.3 | ns |
| **Height (cm)** | 173.1 ± 2.8 | 171.2 ± 2.5 | 170.6 ± 2.4 | ns |
| **Weight (kg)** | 70.6 ± 2.8 | 125.2 ± 5.3 | 93.6 ± 5.2 | *P=0.0002 |
| **BMI (kg/m<sup>2</sup>)** | 23.4 ± 0.4 | 42.5 ± 0.9 | 32.0 ± 1.3 | *P<0.0001 |
| **Waist (cm)** | 78 ± 2 | 123.9 ± 2.2 | 98 ± 5 | *P=0.0002 |
| **Triglycerides**<br>**(mg/dl)** | 0.7 ± 0.07 | 1.1 ± 0.1 | 0.7 ± 0.06 | *P=0.049 |
| **Glucose**<br>**(mmol/l)** | 5.1 ± 0.1 | 5.9 ± 0.1 | 4.8 ± 0.1 | *P<0.0001 |
| **HbA1c**<br>**(mmol/mol)** | 32.5 ± 0.7 | 35.4 ± 1.1 | 32.9 ± 0.9 | *P=0.0105 |
| **Insulin**<br>**sensitivity index**<br>**(M-value)** | 11.9 ± 1.0 | 5.6 ± 0.6 | 7.9 ± 0.5 | *P=0.0018 |

Data is represented as Mean ± SEM. \*Paired t-tests represent differences obese baseline vs obese post bariatric surgery. ns=not significant.

